# Reliability of resting-state functional connectivity in the human spinal cord: assessing the impact of distinct noise sources

**DOI:** 10.1101/2022.12.23.521768

**Authors:** Merve Kaptan, Ulrike Horn, S. Johanna Vannesjo, Toralf Mildner, Nikolaus Weiskopf, Jürgen Finsterbusch, Jonathan C.W. Brooks, Falk Eippert

## Abstract

The investigation of spontaneous fluctuations of the blood-oxygen-level-dependent (BOLD) signal has recently been extended from the brain to the spinal cord, where it has stimulated interest from a clinical perspective. A number of resting-state functional magnetic resonance imaging (fMRI) studies have demonstrated robust functional connectivity between the time series of BOLD fluctuations in bilateral dorsal horns and between those in bilateral ventral horns, in line with the functional neuroanatomy of the spinal cord. A necessary step prior to extension to clinical studies is assessing the reliability of such resting-state signals, which we aimed to do here in a group of 45 healthy young adults at the clinically prevalent field strength of 3T. When investigating connectivity in the entire cervical spinal cord, we observed fair to good reliability for dorsal-dorsal and ventral-ventral connectivity, whereas reliability was poor for within- and between-hemicord dorsal-ventral connectivity. Considering how prone spinal cord fMRI is to noise, we extensively investigated the impact of distinct noise sources and made two crucial observations: removal of physiological noise led to a reduction in functional connectivity strength and reliability – due to the removal of stable and participant-specific noise patterns – whereas removal of thermal noise considerably increased the detectability of functional connectivity without a clear influence on reliability. Finally, we also assessed connectivity within spinal cord segments and observed that while the pattern of connectivity was similar to that of whole cervical cord, reliability at the level of single segments was consistently poor. Taken together, our results demonstrate the presence of reliable resting-state functional connectivity in the human spinal cord even after thoroughly accounting for physiological and thermal noise, but at the same time urge caution if focal changes in connectivity (e.g. due to segmental lesions) are to be studied, especially in a longitudinal manner.

## 1. Introduction

Over the last decades, the spatiotemporal organization of spontaneous fluctuations of BOLD signals in the brain has been widely investigated and intrinsic resting-state networks have been considered as building blocks of brain function that are relevant for cognition and behavior (Deco et al., 2011; Fox & Raichle, 2007; Petersen & Sporns, 2015; Raichle et al., 2001; Wig, 2017). With a delay of about 20 years and on a much smaller scale, a similar perspective has opened up for spinal cord function, with resting-state fMRI studies demonstrating that spontaneous BOLD fluctuations of the spinal cord are spatiotemporally organized as well (Barry et al., 2014; Barry et al., 2016; Barry et al., 2018; Conrad et al., 2018; Eippert et al., 2017a; Harita & Stroman, 2017; Harita et al., 2018; Hu et al., 2018; Ioachim et al., 2019; Ioachim et al., 2020; Kinany et al., 2020; Kong et al., 2014; Liu et al., 2016a; Liu et al., 2016b; Martucci et al., 2019; Martucci et al., 2021; San Emeterio Nateras et al., 2016; Vahdat et al., 2020; Weber et al., 2018; Wei et al., 2009; for a review see Harrison et al., 2021). More specifically, region-of-interest (ROI) based functional connectivity techniques have revealed statistically significant connectivity between the time series of bilateral ventral horns as well as between bilateral dorsal horns in humans and similar functional connectivity patterns have been identified in non-human primates and rodents as well (Chen et al., 2015; Wu et al., 2018; Wu et al., 2019). Since the dorsal horns receive somatosensory information from the body and the ventral horns contain cell bodies of the motor neurons (Hochman, 2007), the observed connectivity patterns appear to be well aligned with the spinal cord’s functional organization.

Resting-state fMRI metrics are often considered in the context of biomarker development (Hohenfeld et al., 2018; Parkes et al., 2018; Pfannmöller & Lotze, 2019), i.e. for monitoring and prediction of disease progression or treatment response. This approach could obviously be extended towards the spinal cord as well (e.g. in the context of recovery after spinal cord injury) and first steps have already been taken in this direction by assessing changes in spinal cord resting-state connectivity in sensory and motor disorders with diffuse or localized spinal pathology (Chen et al., 2015; Combes et al., 2022; Conrad et al., 2018; Martucci et al., 2019). However, before the clinical utility of resting-state metrics can be established, a necessary first step is to assess their reliability as well as the factors that influence it. In this respect, it is important to note that only a very limited number of studies have investigated the test-retest reliability (i.e., the stability of a measure under repeated measures; Shrout and Fleiss, 1979; Shrout and Lane, 2012) of resting-state networks in the human spinal cord: only one study at 7T (Barry et al. 2016) and five studies at the clinically-relevant field strength of 3T (Barry et al., 2018; Hu et al., 2018; Kowalczyk et al., 2023; Liu et al., 2016; San Emeterio Nateras et al., 2016), though these latter ones except Kowalczyk et al. (2023) had rather small sample sizes (N=1 and N=10).

These studies provided an initial assessment of test-retest reliability, but did not investigate the factors that might shape reliability in-depth. Given the susceptibility of spinal cord fMRI to the detrimental influence of noise (Cohen-Adad et al., 2010; for review, see Fratini et al., 2014; Eippert et al., 2017b), it is however essential to understand how distinct noise sources might impact spinal cord resting-state functional connectivity and its reliability – a relationship that, even in the brain, is not necessarily straightforward (Birn et al., 2014; Noble et al., 2019; Shirer et al., 2015). A first noise source of relevance is physiological noise of cardiac and respiratory origin, to which spinal cord fMRI is especially prone (Harita & Stroman, 2017; Piché et al., 2009; Verma & Cohen-Adad, 2014). Physiological noise of structured nature is particularly detrimental for resting-state fMRI studies as one cannot explicitly model the intrinsic activity of interest (unlike in task-based fMRI), which makes it more challenging to attribute the observed results to the underlying neuronal activity instead of non-neural confounds (Birn, 2012; Birn et al., 2014; Murphy et al., 2013). Another major source of noise that influences fMRI measurements is thermal noise (Edelstein et al., 1986; Hoult & Richards, 1976), which has not been investigated in the context of spinal cord fMRI to our knowledge. While thermal noise – whose principal source is the thermal fluctuations within the subject that is imaged, followed by noise due to scanner electronics – is not structured, its removal may further benefit the detectability of BOLD signals of interest (Ades-Aron et al., 2021a; Adhikari et al., 2019; Dowdle et al., 2023; Vizioli et al., 2021).

Considering all the above, the aims of the current study are as follows. First, we aim to replicate previous resting-state fMRI functional connectivity results and assess their test-retest reliability in a large sample (N=45) at the clinically-relevant field strength of 3T across the entire cervical spinal cord. Second, we aim to assess how structured (physiological) and unstructured (thermal) noise sources impact functional connectivity and its reliability. Finally, we aim to investigate more localized aspects of functional connectivity and its reliability, namely within each spinal cord segment, i.e. the macro-scale building blocks of spinal cord organization.

## 2. Methods

### 2.1. Participants

This study is based on the participant sample of Kaptan et al. (2022), which contained data from 48 healthy participants. As our focus in the current study was on assessing the influence of different noise sources on the reliability of resting-state functional connectivity, data from three participants had to be discarded due to technical problems in the acquisition of peripheral physiological data (i.e., corrupted ECG-recordings), thus leading to a final sample size of 45 participants (20 females, age: 27 ± 3.8). All participants provided written informed consent and the study was approved by the Ethics Committee at the Medical Faculty of the University of Leipzig.

### 2.2. Data acquisition

All measurements were performed on a 3T whole-body Siemens Prisma MRI System (Siemens, Erlangen, Germany) equipped with a whole-body radio-frequency (RF) transmit coil, a 64-channel RF head-and-neck coil, and a 32-channel RF spine-array, using the head coil element groups 5–7, the neck coil element groups 1 and 2, and spine coil element group 1 (all receive-only). Before the start of data acquisition, typical instructions for spinal MRI studies were given to the participants (i.e., they were told not to move, to avoid excessive swallowing and to breathe normally; see Cohen-Adad et al., 2021). The here-described data are part of a larger methodological project: we thus only describe the relevant parts – two functional acquisitions and one structural acquisition – and refer the interested reader to the methodological publication for further details on this dataset (Kaptan et al., 2022).

Functional runs consisted of 250 single-shot 2D gradient-echo EPI volumes (acquisition time: 578s) that covered the spinal cord from the 2nd cervical vertebra to the 1st thoracic vertebra and were acquired with the following parameters: slice orientation: transverse oblique; number of slices: 24; slice thickness: 5.0mm; field of view: 128×128mm^2^, in-plane resolution: 1.0 × 1.0mm^2^; TR: 2312ms; TE: 40ms; excitation flip angle: 84°, GRAPPA acceleration factor: 2; partial Fourier factor: 7/8; phase-encoding direction: anterior-to-posterior; echo spacing: 0.93ms; bandwidth per pixel: 1220 Hz/Pixel. Both functional runs employed slice-specific z-shimming (Finsterbusch et al., 2012) in order to overcome the signal-loss that occurs due to local magnetic field inhomogeneities. The two runs only differed according to the selection method of slice-specific z-shims: this occurred either manually or automatically (Kaptan et al., 2022). The two runs were separated from each other by a maximum of ∼10 minutes, did not show a systematic order difference (the run with manual selection of z-shims occurred before the run with automatic selection of z-shims in 23 of the 45 participants) and exhibited highly similar gray matter tSNR (run with manual selection of z-shims: 15.7 ± 1.3; run with automatic selection of z-shims: 15.4 ± 1.3; mean ± standard deviation; see also Figure S1 for voxel-wise gray matter tSNR maps). During each of the runs, participants were presented with a white cross-hair on a gray background, which they were asked to fixate on.

Additionally, a high-resolution T2-weighted acquisition (3D sagittal SPACE sequence, Cohen-Adad et al., 2021; 64 sagittal slices; resolution: 0.8×0.8×0.8mm^3^; field-of-view: 256×256mm^2^; TE: 120ms; flip angle: 120°; TR: 1500ms; GRAPPA acceleration factor: 3; acquisition time: 4.02min) was obtained for registration purposes.

During fMRI data acquisition, we also acquired peripheral physiological signals in order to perform physiological noise modelling: respiratory data were acquired via a breathing belt and cardiac data were acquired via ECG electrodes (BrainAmp ExG system; Brain Products GmbH, Gilching, Germany). Data acquisition occurred with a sampling-rate of 1kHz and included scanner triggers to allow for synchronization of data streams.

### 2.3. Data preprocessing

Preprocessing steps were performed using MATLAB (version 2021a), EEGLAB (version 2019.0; Delorme & Makeig, 2004), FMRIB Software Library (FSL; version 6.0.3; Jenkinson et al., 2012), and Spinal Cord Toolbox (SCT; version 4.2.2; De Leener et al., 2017).

#### 2.3.1. Preprocessing of physiological data

ECG data were processed within EEGLAB (Delorme & Makeig, 2004) using the FMRIB plug-in (https://fsl.fmrib.ox.ac.uk/eeglab/fmribplugin/). This algorithm allows for the correction of gradient artifacts in the ECG signal caused by the switching of magnetic gradients during fMRI acquisitions (Niazy et al., 2005). R-peaks were automatically detected after correction and where necessary manual corrections were carried out using in-house MATLAB scripts.

We calculated the heart-period (i.e., R-R interval) in milliseconds as the average difference in time between each R peak for each functional run. In addition to that, we assessed heart-period variability by calculating the standard deviation of R-R intervals (Shaffer & Ginsberg, 2017) within each of the two functional runs.

The respiratory period was calculated as described by Bach and colleagues (2016). More specifically, the respiration traces were i) mean-centered, ii) filtered with a band pass filter (cut-off frequencies: 0.01 Hz and 0.6 Hz), and iii) median filtered over 1s. The start of inspiration was defined as a negative zero-crossing. After each detected cycle, a 1s refractory period was imposed, to account for residual signal noise that may lead to the occurrence of several zero-crossings on the same respiratory cycle (Bach et al., 2016). We report the mean and standard deviation of the respiratory period in seconds.

#### 2.3.2. Preprocessing of fMRI data

##### 2.3.2.1. Motion-correction

For each functional run, a slice-wise motion correction procedure with regularization in z-direction (as implemented in SCT, “*sct_fmri_moco*”) was employed in two steps. First, the 250 volumes of each run were averaged to create a mean image, and this mean image was used to automatically determine the centerline of the cord. A cylindrical mask (with a diameter of 41mm) was generated based on this centerline and used during the motion-correction procedure to ensure that regions moving independently from the cord would not adversely impact the motion-correction. The previously-created mean image was used as a target for the first iteration of slice-wise motion correction with a 2^nd^ degree polynomial and spline interpolation. In the second step, the mean of motion-corrected time series from the first step served as a target image for the second iteration of motion-correction, which was applied to the raw images (with the same algorithm parameters).

##### 2.3.2.2. Segmentation

For the functional runs, binary masks/segmentations of the spinal cord were manually created based on each mean image after motion-correction. We employed a manual segmentation instead of an automated segmentation to ensure that the segmentation quality did not adversely affect the registration procedure (see below), which was dependent on the segmentation.

Binary masks/segmentations of the spinal cord obtained from the T2-weighted images were created automatically using the ‘*sct_deepseg*’ approach of SCT (Gros et al., 2019).

##### 2.3.2.3. Registration

Functional connectivity analyses were performed in native space to make them comparable to those of a previous study on resting-state functional connectivity and its reliability by Barry and colleagues (2016). However, a registration procedure to the PAM50 template space (De Leener et al., 2018) was still performed in order to obtain the warping fields that allowed to bring region-specific probabilistic masks from PAM50 template space to each individual’s native space (‘*sct_warp_template’*).

First, anatomical T2-weighted images were normalized to the template space with the following three consecutive steps (*’sct_register_to_template’*): i) the spinal cord was straightened using the binary cord segmentation, ii) the automatically labelled C2-C7 vertebral levels (created via ‘*sct_label_vertebrae*’, with manual corrections when deemed necessary) were used for the vertebral alignment between the template and the anatomical images, iii) the anatomical images were registered to the template using non-rigid segmentation-based transformations.

Second, the T2-weighted PAM50 template was registered to the mean of motion-corrected functional images using non-rigid transformations (‘*sct_register_multimodal*’; with the initial step using the inverse warping field obtained from the registration of the T2-weighted anatomical image to the template image). The resulting warping fields obtained from this registration were then applied to the PAM50 probabilistic gray matter and segmental level masks to bring them into the native space where connectivity estimation and statistical analyses were carried out.

#### 2.3.3. Denoising

As we aimed to investigate the effect of various noise sources on resting-state functional connectivity and its reliability, we employed different denoising pipelines to assess the impact of specific noise sources.

##### 2.3.3.1. Physiological noise

First, we employed a processing pipeline that does not explicitly account for any specific noise source – from now on we refer to this pipeline as ‘*baseline*’ throughout the manuscript. The baseline denoising pipeline consisted of i) motion-correction, ii) high-pass filtering (with a 100s cut-off), and iii) “motion-censoring”. Censoring was necessary to ensure that outlier volumes that were either inadvertently introduced by the motion-correction algorithm or that occurred due to a sudden large movement of participants did not artificially inflate the connectivity estimates (as outlier volumes can create spikes in the signal time series of ROIs). The outlier volumes were determined using the dVARS (the root mean square difference between successive volumes; Smyser et al., 2011) and refRMS (root mean square intensity difference of each volume to the reference volume) metrics as implemented in the ‘*fsl_motion_outliers’* function of FSL. Volumes presenting with dVARS or refRMS values two standard deviations above the mean values of each run were selected as outliers. In the later occurring GLM estimation, these outlier volumes were modelled as individual regressors (on average, 4.67 ± 3.15 volumes were identified as outliers across all participants and sessions, i.e. less than 2% of the volumes).

Second, physiological noise modelling (PNM; Brooks et al., 2008) was used to obtain slice-specific regressors to account for physiological confounds. PNM is a modification of the RETROICOR approach (Glover et al., 2000) and creates slice-specific regressors via calculating the cardiac and respiratory phase of each slice by modelling them via Fourier basis series with a combination of sine and cosine harmonics (Brooks et al., 2008; Kong et al., 2012). We utilized regressors up to the fourth harmonic – resulting in a total of 16 regressors – to account for cardiac and respiratory processes, and another 16 regressors to account for their interactions, resulting in a total of 32 regressors (Brooks et al., 2008; Kong et al., 2012). In addition to that, a slice-specific CSF regressor was created (as implemented in PNM) by extracting the signal from the voxels whose variance were in the top 10 percentile within a region including both the spinal cord and CSF space. In post-hoc analyses, we also created slice-specific white-matter (WM) regressors in the following way: we i) registered the PAM50 WM template to native space, ii) subtracted the native space unthresholded gray matter template (in order to prevent overlap with the gray matter mask) and ii) obtained the average time series from the resulting mask (in order to be used as WM regressor). Note that all noise regressors were high-pass filtered with the same 100s cut-off prior to noise regression to prevent spectral misspecification (Hallquist et al., 2013).

Third, a specific set of regressors that account for different physiological noise sources was then added to the baseline denoising pipeline, and regressed out from the functional data using FEAT (FMRI Expert Analysis Tool; http://fsl.fmrib.ox.ac.uk/fsl/fslwiki/FEAT), resulting in the seven different denoising pipelines listed below:

i. Baseline (consisting of motion-correction, high-pass filtering and censoring)
ii. Baseline + slice-specific motion-correction estimates (x- and y-translation; automatically obtained from the slice-wise motion correction procedure)
iii. Baseline + CSF signal
iv. Baseline + eight respiratory regressors
v. Baseline + eight cardiac regressors
vi. Baseline + thirty-two PNM regressors (including eight respiratory regressors, eight cardiac regressors, and 16 interaction regressors)
vii. Maximal (motion-correction, high-pass filtering, censoring, slice-specific motion correction regressors, 32 PNM regressors and a CSF regressor)

The residuals obtained from each of the denoising pipelines were then used for further analysis. Please note that while we did not include a pre-whitening step in our above-mentioned denoising pipelines, we assessed the impact of pre-whitening carried out using FILM (FMRIB’s Improved Linear Model with local autocorrelation correction; Woolrich et al., 2001) by comparing maximal denoising with maximal denoising + FILM pre-whitening (see Table S1). In post-hoc analyses, we also assessed the impact of WM regression by adding a WM regressor to baseline processing, as well as to maximal processing (see Table S2).

##### 2.3.3.2. Thermal noise

Another major source of noise that contributes to the variability of fMRI time series is zero-mean Gaussian thermal noise which arises from thermal fluctuations within the participant, as well as scanner electronics (Edelstein et al., 1986; Hoult & Richards, 1976). Here, we employed two different approaches to address the influence of thermal noise: spatial smoothing and denoising based on Marchenko-Pastur Principle Component Analysis (MP-PCA; Marčenko & Pastur, 1967; Veraart et al., 2016a; Veraart et al., 2016b), either of which was employed before GLM-based physiological noise correction via the maximal denoising pipeline was carried out. Spatial smoothing was implemented in FEAT with isotropic Gaussian kernels of either 2mm or 4mm FWHM. Non-local MP-PCA was implemented using an openly available MATLAB algorithm (http://github.com/NYU-DiffusionMRI/mppca_denoise; Ades-Aron et al., 2021b) and was applied to the entire fMRI time series data (dimensions [x, y, z, time]: 128 × 128 × 24 × 250) before motion correction. In the context of MRI, MP-PCA was originally evaluated for thermal noise reduction in diffusion MRI data (Veraart et al., 2016a; Veraart et al., 2016b), but has recently also been applied to task-based (Ades-Aron et al., 2021a) and resting-state (Adhikari et al., 2019) fMRI data of the brain, aiming to minimize the contributions of thermal noise to fMRI time series without altering the spatial resolution.

Finally, in order to estimate the effect of thermal noise removal – via smoothing or MP-PCA – on the data’s spatial smoothness, we estimated the spatial autocorrelation function of the residuals within the spinal cord after each of four processing pipelines (maximal, maximal + MP-PCA, maximal + smoothing 2mm, maximal + smoothing 4mm) using the 3dFWHMx function of AFNI (Cox et al., 2017). The smoothness estimates were derived from AFNI’s mixed gaussian and mono-exponential decay model and we report the effective (combined) smoothness value after each denoising approach (already incorporating smoothness changes introduced during motion correction).

### 2.4. Statistical analysis

#### 2.4.1. Functional connectivity calculation

Functional connectivity was assessed using an ROI-based approach. The ROI masks were created using the probabilistic PAM50 gray matter masks that were warped from template space to the native space of each participant (see section 2.3.2.3). In native space, the probabilistic gray matter masks were thresholded at 70% for each slice separately to ensure that there were no voxels shared between distinct ROIs. Within a slice, the ROIs typically contained 1.6 and 1.5 voxels in the left and right dorsal horns, and 1.9 and 1.9 voxels in the left and right ventral horns, respectively (average over slices and participants). *Slice-specific* time courses were then extracted via averaging the signal over the voxels within each of the four ROIs (left dorsal horn, left ventral horn, right dorsal horn, and right ventral horn).

Next, slice-wise correlations between ROIs were calculated using the Pearson correlation coefficient (see Supplementary Material for an explanation as to why correlations were calculated slice-wise). In order to address the effects of any remaining signal fluctuations that might be shared between the ROIs (e.g. residual movement or physiological noise effects) we also calculated slice-wise partial correlation coefficients (Figure S2): for instance, to calculate the partial correlation on a given slice between time series from left and right dorsal horn, the time series from left and right ventral horn of that slice were used as controlling variables. The dorsal-ventral correlations within each hemicord (left dorsal with left ventral and right dorsal with right ventral), as well as between hemicords (left dorsal with right ventral and right dorsal with left ventral) were averaged, yielding one within-hemicord and one between-hemicord dorsal-ventral connectivity value for each participant (similar to Eippert et al. 2017a, who did not observe any significant laterality differences). The slice-wise correlation coefficients were then averaged over all slices along the superior-inferior axis of the cord, yielding four functional connectivity estimates for each participant: dorsal-dorsal, ventral-ventral, dorsal-ventral within-hemicord and dorsal-ventral between-hemicord. This averaging of correlation values might lead to a slight conservative bias in our results as we did not perform Fischer’s z-transformation prior to averaging, however, this is assumed to be negligible (Silver & Dunlap, 1987; Corey et al., 1998; Eippert et al., 2017a). Note that only those slices that were assigned to C3-T1 probabilistic segmental levels were included, resulting in a variable number of slices across different participants due to the anatomy of the participants (depending on the coverage of the EPI slice-stack during acquisition). At the group-level, we report the mean r value, i.e. averaged across two sessions and averaged across participants.

The significance of the functional connectivity estimates or the difference between them (depending on the aim of the analysis) were assessed using permutation-based tests implemented in the Permutation Analysis of Linear Models software (PALM; Winkler et al., 2014). The number of permutations was set to 10,000 and we report two-tailed family-wise error (FWE) corrected p-values (adjusted according to the number of tests performed).

##### 2.4.1.1. Within-segment functional connectivity

In order to provide insights into the segment-wise organization of functional connectivity, we also investigated the functional connectivity within each spinal segment covered by our imaging volume; those included all segments between the third cervical (C3) and first thoracic segment (T1). Therefore, probabilistic segmental levels from PAM50 template space were first warped to each participant’s native space (see section 2.3.2.3). Then, to guarantee that there was no overlap between neighboring segments, the slice with the highest probability of belonging to a specific segmental level and the slice above and below were assigned to the corresponding segment. This procedure ensured that there were a similar number of slices for each segment and led to a 15 mm segment length, which is in line with empirical measurements of cervical segment length based on post-mortem data (Ko et al., 2004). Slice-wise functional connectivity was calculated as described above and the correlation values for slices within each segment were averaged. The connectivity strength for each segment was tested against 0 via permutation tests as described above (see section 2.4.1). Please note that for all within-segment analyses, we used data that had undergone the maximal denoising pipeline for physiological noise correction and were also corrected for thermal noise via MP-PCA, as our whole-cord analyses had suggested that this was the optimal processing pipeline.

#### 2.4.2. tSNR and explained variance

In order to provide further insights into the effects of the removal of various noise sources, we also calculated the gray matter temporal signal-to-noise ratio (tSNR) and the explained variance of the gray matter time series *for each denoising step* (please note that motion correction, high-pass filtering and motion-censoring was always performed). Voxelwise gray matter tSNR values were calculated for each functional run via dividing each voxel’s temporal mean by its temporal standard deviation (Parrish et al., 2000). The impact of various noise sources on gray matter tSNR was assessed by comparing the tSNR values obtained after each denoising pipeline to the baseline denoising procedure – in addition to reporting descriptive values (% change) we also employed permutation-based tests as described above (see section 2.4.1) and report FWE-corrected p-values. Following Birn et al. (2014), the variance of gray matter time series explained by each denoising pipeline (R^2^) was calculated by computing the fractional reduction in signal variance. tSNR and explained variance for each gray matter region were extracted using the native-space thresholded and binarized PAM50 gray matter masks that were also used to calculate functional connectivity.

#### 2.4.3. Estimation of reliability

The central aspect of this manuscript concerns the reliability of resting-state functional connectivity in the human spinal cord. While different fields have come to rely on different operationalizations of reliability (for an in-depth discussion, see Brandmaier et al., 2018), we here follow the tradition in resting-state functional connectivity research and employ the intra-class correlation coefficient (ICC) for assessing reliability (see also Noble et al., 2020). Considering that spinal cord fMRI is severely impacted by different noise sources, our reliability investigation was not only focused on the connectivity metrics, but also possibly contributing factors. Thus, we calculated the test-retest reliability for each of the following aspects: i) functional connectivity, ii) tSNR, iii) motion metrics (DVARS, refRMS), iv) cardiac metrics (mean heart period, heart period variability), v) respiratory metrics (mean respiratory period, respiratory period variability), and vi) explained variance of gray matter time series.

For each of these metrics, we first created a 45×2 (i.e. participants×sessions) matrix and then assessed the reliability using the ‘Case 2’ intraclass correlation coefficient (ICC(2,1); two-way random effects model; McGraw & Wong, 1996; Shrout & Fleiss, 1979); this is often also referred to as ‘absolute agreement’ (Molloy & Birn, 2014). ICC(2,1) is defined as the following:

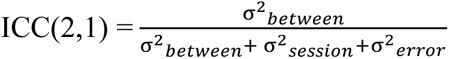

Where 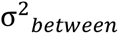 corresponds to the variance among persons (between participant) and 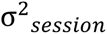 corresponds to the variance between sessions. Given its formula, the ICC shows what proportion of the total variance can be attributed to between-persons differences (Brandmaier et al., 2018; Noble et al., 2019).

We also aimed to provide an estimate of uncertainty, and thus calculated the 95% confidence interval (CI) of ICC values via non-parametric bootstrapping performed in MATLAB. Throughout the manuscript, ICC values are interpreted according to standard procedures: poor <0.4, fair 0.4–0.59, good 0.6–0.74, excellent ≥0.75 (Cicchetti & Sparrow, 1981; Hallgren, 2012).

### 2.5. Open science statement

All the code necessary to reproduce the reported results is available on GitHub (https://github.com/eippertlab/restingstate-reliability-spinalcord). The underlying data are available in BIDS-format via OpenNeuro (https://openneuro.org/datasets/ds004386; note that the dataset is currently only accessible to reviewers). The intended data-sharing via OpenNeuro was mentioned in the Informed Consent Form signed by the participants and approved by the Ethics Committee at the Medical Faculty of the University of Leipzig.

## 3. Results

### 3.1. Replication and extension of previous resting-state functional connectivity results

Our first aim was to i) replicate previous ROI-based resting-state functional connectivity fMRI findings and ii) quantify the test-retest reliability of resting-state functional connectivity at 3T in human spinal cord. To this end, we assessed connectivity between the dorsal horns, between the ventral horns and between the within-hemicord dorsal and ventral horns as well as between-hemicord dorsal and ventral horns (Figure 1A). All connectivity estimations were carried out on data that were subjected to extensive correction for physiological noise (i.e. the ‘maximal’ denoising pipeline), as is typical in spinal fMRI. To control for non-specific factors, we explored tSNR differences between the different horns, but observed rather similar group-averaged gray matter tSNR (even though the tSNR of ventral horns were slightly higher (6.8%) compared to the dorsal horns), with the range of variation across participants also being similar (Figure 1B).

**Figure 1.**
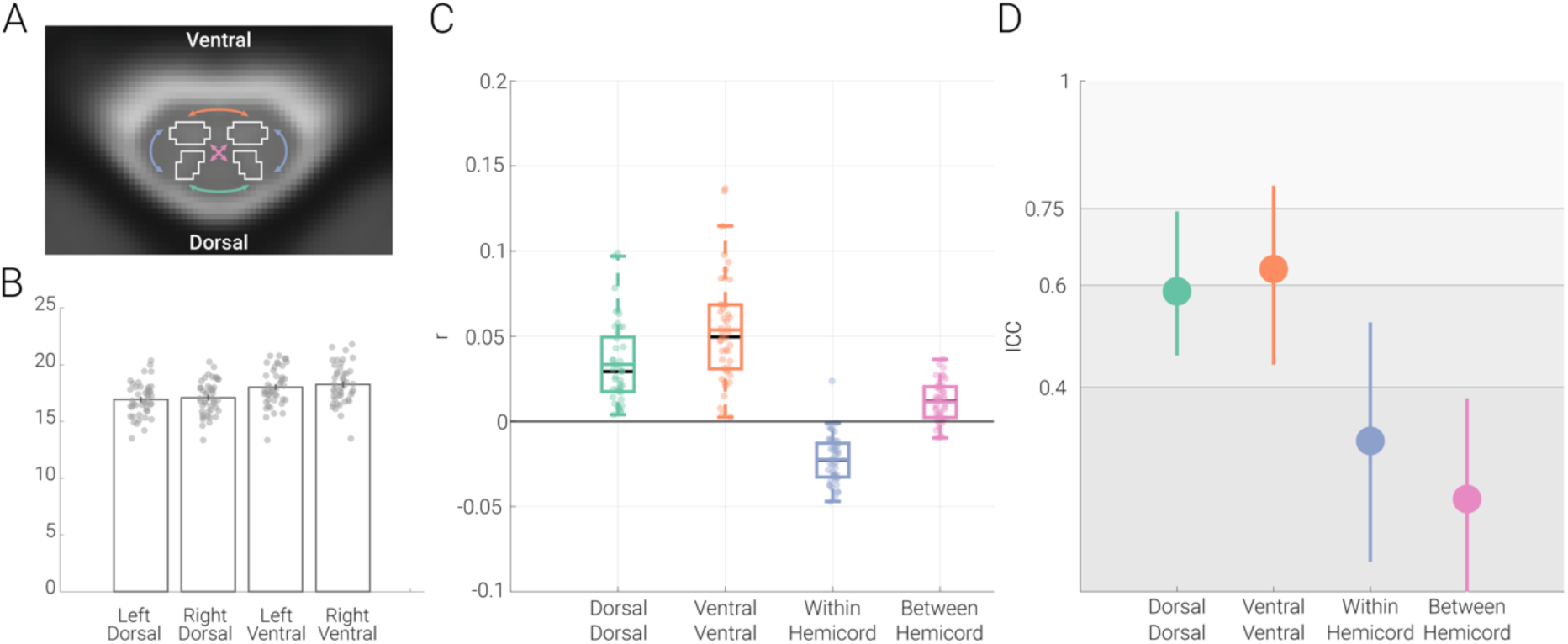
Resting-state functional connectivity and its reliability. **A. Functional connectivity calculation.** An exemplary transverse slice taken from the T2*-weighted PAM50 template (at segmental level C6) is shown with the gray matter masks overlaid as contours; please note that while for visualization purposes we display an exemplary slice from the PAM50 template here, all analyses were carried out in native space. The coloured arrows indicate the four different types of ROI-to-ROI connectivity that we investigated: dorsal-dorsal in green, ventral-ventral in orange, within-hemicord dorsal-ventral in blue, and between-hemicord dorsal-ventral in pink. **B. Gray matter tSNR.** Bar graphs show the tSNR for each of the gray matter ROIs. The vertical lines on the bars depict the standard error of the mean and the circles indicate participant-specific values. **C. Resting-state functional connectivity of the cervical cord.** Pearson correlation values (averaged across two sessions) between the time-courses of different ROIs are shown with box plots. For the box plots, the median is denoted by the black central line and the mean is denoted by the colored central line. The boxes represent the interquartile range and the whiskers encompass ∼99% of the data. Correlation values from individual participants are shown with circles. **D. Test-retest reliability of resting-state connectivity.** ICC values for each connection are indicated via the circles, with the vertical lines representing the 95% confidence intervals. The gray scale background reflects the ICC ranges (as defined by Cicchetti & Sparrow (1981) and (Hallgren, 2012)): poor <0.4, fair 0.4–0.59, good 0.6–0.74, excellent ≥0.75.

We observed highly significant positive connectivity between the dorsal horns (r = 0.03; t = 9.5; p < 0.001) as well as between the ventral horns (r = 0.05; t = 11.6; p < 0.001) and were thus able to replicate previous findings. Additionally, we observed significant negative dorsal-ventral connectivity within hemicords (r = -0.02; t = -10.7; p < 0.001) and positive dorsal-ventral connectivity between hemicords (r = 0.01; t = 6.7; p < 0.001), but these were weaker than the dorsal and ventral connectivity (Figure 1C). With regards to the robustness of these results at the individual level, 100% of the participants exhibited positive dorsal-dorsal and ventral-ventral connectivity, while 98% of participants exhibited negative dorsal-ventral within-hemicord connectivity and 84% of participants demonstrated positive dorsal-ventral between-hemicord connectivity.

In terms of the reliability of these connectivity patterns, the ICC of dorsal-dorsal connectivity (0.59, CI: 0.46 – 0.74) and of ventral-ventral connectivity (0.63, CI: 0.44 – 0.79) was in the upper part of the fair and the lower part of the good range, respectively, whereas the reliability of within- and between-hemicord dorsal-ventral connectivity was clearly in the poor range (within-hemicord: 0.30, CI: 0.06 – 0.53 ; between-hemicord: 0.18, CI : -0.03 – 0.38; Figure 1D). Both connectivity amplitude and reliability were also assessed by i) replacing Pearson correlation with partial correlation (in order to account for the effects of any possibly remaining global signal fluctuations) and ii) adding a pre-whitening step during the GLM estimation (in order to account for the temporal autocorrelation of the BOLD data), but neither of these approaches led to a relevant change in the here-reported results (see Figure S2 and Table S1, respectively).

### 3.2. Impact of noise sources on resting-state functional connectivity and its reliability

Considering that spinal cord fMRI is severely signal-to-noise limited due to the impact of various noise sources, we next investigated the relevance of each of these noise sources for the estimation of functional connectivity and its reliability. While the above-reported results were obtained after typical physiological noise correction procedures, we now separately assess physiological noise sources as well as thermal noise, which has hitherto been neglected in spinal cord fMRI. The effects of each noise source were evaluated by assessing the change in connectivity amplitude and reliability after it was removed.

#### 3.2.1. Physiological noise and amplitude of functional connectivity

There are several general observations regarding the effects of physiological noise sources on functional connectivity (Figure 2; Tables 1 & 2). First, no matter which noise source was corrected for, the sign of the correlation stayed the same for all four connections and all four connections remained significant, indicating their robustness. Second, the (relatively weaker) within-hemicord and between-hemicord connectivity strength was not systematically impacted by physiological noise correction. Third, and most importantly, dorsal-dorsal and ventral-ventral connections showed a consistent reduction in connectivity strength with increasingly stringent denoising. This latter point was also evident statistically, where a significant reduction in connectivity strength was observed for all noise sources, which became even more pronounced when combining the different noise regressors into combined sets (e.g. *PNM* pipeline and *maximal* pipeline; see Table 1). Interestingly, despite the strong reduction in correlation amplitude for dorsal-dorsal and ventral-ventral connections (of at least 50%) from the *baseline* to the *maximal* pipeline, the results remained clearly significant in the latter, which was likely due to the reduction in the inter-individual spread of amplitudes (i.e. higher precision). Supporting this overall pattern, highly similar results were obtained when Pearson correlation was replaced by partial correlation (Figure S2); post-hoc analyses including white-matter regression did not lead to meaningful changes (Table S2).

**Figure 2.**
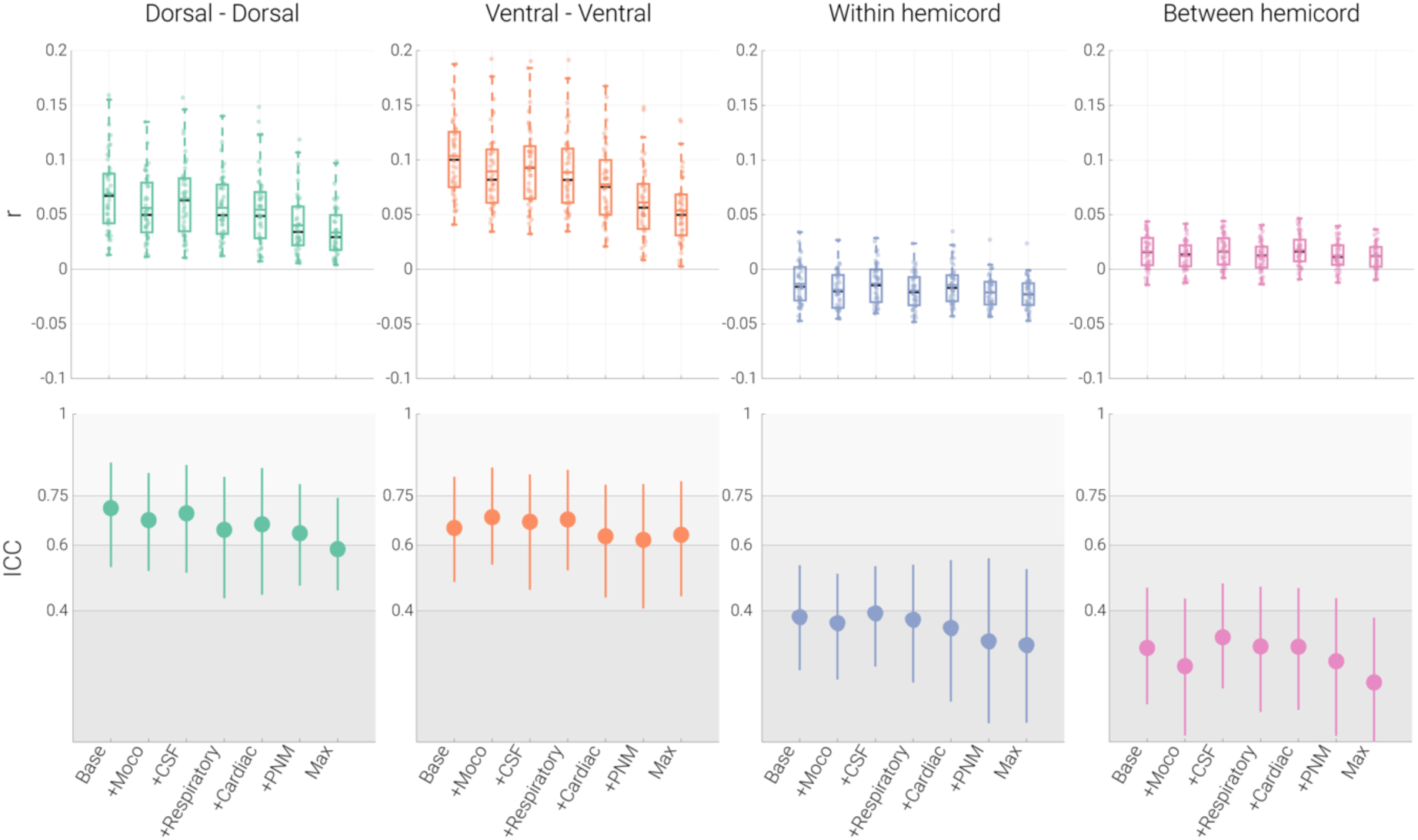
Effects of physiological noise. The top panel depicts Pearson correlation values (averaged within a participant across the two runs) between the time-courses of different ROIs via box plots for the seven denoising pipelines (Base: baseline processing; +Moco: baseline + slice-specific motion-correction estimates; +CSF: baseline + CSF signal; +Respiratory: baseline + eight respiratory regressors; +cardiac: baseline + eight cardiac regressors; +PNM: baseline + thirty-two PNM regressors; Max: baseline processing, slice-specific motion correction estimates, 32 PNM regressors and a CSF regressor). For the box plots, the median and mean are denoted by black and colored central lines, respectively. The boxes represent the interquartile range, with the whiskers encompassing ∼99% of the data and the circles representing individual participants. The bottom panel depicts ICC values for each the different pipelines via the circles, with the vertical lines representing the 95% confidence intervals. The gray scale background reflects the ICC ranges (as defined by Cicchetti & Sparrow (1981) and Hallgren (2012)): poor <0.4, fair 0.4–0.59, good 0.6–0.74, excellent ≥0.75.

**Table 1.** Functional connectivity and reliability after physiological noise correction. This table depicts functional connectivity and reliability results of each connection across seven denoising pipelines. r represents the mean Pearson correlation across participants, and t and p represent the t-value and two-tailed FWE-corrected (for seven tests) p-value from a permutation test (against 0), respectively. ICC(95% CI) represents ICC(2,1) values and 95% bootstrapped confidence intervals.

|  |  | **Dorsal**<br>**Dorsal** | **Ventral**<br>**Ventral** | **Within**<br>**Hemicord** | **Between**<br>**Hemicord** |
| --- | --- | --- | --- | --- | --- |
| **Baseline** |  | r = 0.07<br>t = 12.9<br>p < 0.001<br>ICC (95% CI) =<br>0.71 (0.53 – 0.85) | r = 0.10<br>t = 18.7<br>p < 0.001<br>ICC (95% CI) =<br>0.65 (0.49 – 0.81) | r = -0.01<br>t = -4.7<br>p < 0.001<br>ICC (95% CI) =<br>0.38 (0.22 – 0.54) | r = 0.02<br>t = 7.1<br>p < 0.001<br>ICC (95% CI) =<br>0.28 (0.11 – 0.47) |
| **Baseline**<br>**Motion**<br>**parameters** | + | r = 0.06<br>t = 12.4<br>p < 0.001<br>ICC (95% CI) =<br>0.68 (0.52 – 0.82) | r = 0.09<br>t = 17.4<br>p < 0.001<br>ICC (95% CI) =<br>0.69 (0.54 – 0.84) | r = -0.02<br>t = -4.7<br>p < 0.001<br>ICC (95% CI) =<br>0.36 (0.19 – 0.51) | r = 0.01<br>t = 7.7<br>p < 0.001<br>ICC (95% CI) =<br>0.23 (0.02 – 0.44) |
| **Baseline + CSF** |  | r = 0.06<br>t = 12.9<br>p < 0.001<br>ICC (95% CI) =<br>0.69 (0.52 – 0.84) | r = 0.09<br>t = 16.9<br>p < 0.001<br>ICC (95% CI) =<br>0.67 (0.46 – 0.82) | r = -0.01<br>t = -7.7<br>p < 0.001<br>ICC (95% CI) =<br>0.39 (0.23 – 0.54) | r = 0.02<br>t = 6.0<br>p < 0.001<br>ICC (95% CI) =<br>0.32 (0.16 – 0.48) |
| **Baseline**<br>**Respiratory** | + | r = 0.06<br>t = 12.8<br>p < 0.001<br>ICC (95% CI) =<br>0.65 (0.44 – 0.81) | r = 0.09<br>t = 16.6<br>p < 0.001<br>ICC (95% CI) =<br>0.68 (0.52 – 0.83) | r = -0.02<br>t = -8.1<br>p < 0.001<br>ICC (95% CI) =<br>0.37 (0.18 – 0.54) | r = 0.01<br>t = 5.7<br>p < 0.001<br>ICC (95% CI) =<br>0.29 (0.09 – 0.47) |
| **Baseline**<br>**Cardiac** | + | r = 0.05<br>t = 10.8<br>p < 0.001<br>ICC (95% CI) =<br>0.66 (0.45 – 0.84) | r = 0.08<br>t = 15.0<br>p < 0.001<br>ICC (95% CI) =<br>0.63 (0.44 – 0.78) | r = -0.01<br>t = -5.5<br>p < 0.001<br>ICC (95% CI) =<br>0.35 (0.12 – 0.55) | r = 0.02<br>t = 8.4<br>p < 0.001<br>ICC (95% CI) =<br>0.29 (0.10 – 0.47) |
| **Baseline**<br>**PNM** | + | r = 0.04<br>t = 9.8<br>p < 0.001<br>ICC (95% CI) =<br>0.64 (0.48 – 0.79) | r = 0.06<br>t = 12.4<br>p < 0.001<br>ICC (95% CI) =<br>0.62 (0.41 – 0.79) | r = -0.02<br>t = -9.4<br>p < 0.001<br>ICC (95% CI) =<br>0.31 (0.06 – 0.56) | r = 0.01<br>t = 6.7<br>p < 0.001<br>ICC (95% CI) =<br>0.25 (0.02 – 0.44) |
| **Maximal** |  | r = 0.03<br>t = 9.5<br>p < 0.001<br>ICC (95% CI) =<br>0.59 (0.46 – 0.74) | r = 0.05<br>t = 11.6<br>p < 0.001<br>ICC (95% CI) =<br>0.63 (0.44 – 0.79) | r = -0.02<br>t = -10.7<br>p < 0.001<br>ICC (95% CI) =<br>0.30 (0.06 – 0.53) | r = 0.01<br>t = 6.7<br>p < 0.001<br>ICC (95% CI) =<br>0.18 (-0.03 – 0.38) |

**Table 2.** Comparison of functional connectivity strength for different denoising pipelines. This table depicts statistical comparisons of the functional connectivity strength (for each of the four connections) for six different denoising pipelines against the baseline pipeline. t and p represent the t-value and two-tailed FWE-corrected (for six tests) p-value from a permutation test against 0 (as values for each connection were subtracted from the baseline functional connectivity values). Note that for within hemicord connectivity (where connectivity values are negative), smaller t-values mean that the negative connectivity gets stronger.

|  |  | **Dorsal**<br>**Dorsal** | **Ventral**<br>**Ventral** | **Within**<br>**Hemicord** | **Between**<br>**Hemicord** |
| --- | --- | --- | --- | --- | --- |
| **Baseline**<br>**Motion**<br>**parameters** | + | t(44) = -8.0<br>p < 0.001 | t(44) = -9.9<br>p < 0.001 | t(44) = -6.1<br>p < 0.001 | t(44) = -5.1<br>p < 0.001 |
| **Baseline**<br>**CSF** | + | t(44) = -6.6<br>p < 0.001 | t(44) = -8.1<br>p < 0.001 | t(44) = 0.4<br>p = 0.98 | t(44) = -0.6<br>p = 0.93 |
| **Baseline**<br>**Respiratory** | + | t(44) = -8.8<br>p < 0.001 | t(44) = -11.3<br>p < 0.001 | t(44) = -7.0<br>p < 0.001 | t(44) = -6.1<br>p < 0.001 |
| **Baseline**<br>**Cardiac** | + | t(44) = -10.5<br>p < 0.001 | t(44) = -11.1<br>p < 0.001 | t(44) = -1.0<br>p = 0.68 | t(44) = 1.1<br>p = 0.64 |
| **Baseline**<br>**PNM** | + | t(44) = -11.4<br>p < 0.001 | t(44) = -16.3<br>p < 0.001 | t(44) = -5.6<br>p < 0.001 | t(44) = -2.6<br>p = 0.04 |
| **Maximal** |  | t(44) = -11.6<br>p < 0.001 | t(44) = -17.8<br>p < 0.001 | t(44) = -5.9<br>p < 0.001 | t(44) = -3.0<br>p = 0.01 |

#### 3.2.2. Physiological noise and reliability of functional connectivity

Similar to the strength of functional connectivity, reliability also decreased with more stringent denoising (Figure 2; Table 1), though now for all four connections: the reliability of dorsal-dorsal connectivity decreased from good to fair (by 17.5%), the reliability of ventral-ventral functional connectivity stayed in the good range with a slight decline (by 3.19%), and the ICC values for within- and between-hemicord connectivity were consistently in the poor range, though with a clear decline of reliability being noticeable (22.5% and 36.7%, respectively). When looking at the influence of single noise sources, it becomes apparent that the strongest drop in reliability is observed due to removal of respiratory noise for dorsal-dorsal connectivity, whereas the removal of cardiac noise leads to the strongest decline of reliability in ventral-ventral connectivity.

The observed decrease in reliability may seem counter-intuitive at first glance, as the removal of physiological noise could be expected to increase reliability. However, such a pattern could arise if i) the noise is spatially structured (which is known to be the case for physiological noise) and ii) the processes that generate noise present with high reliability, which we set out to probe here. We noticed that metrics of motion (DVARS and refRMS), cardiac activity (mean heart period and heart period variability) and respiratory activity (mean respiratory period and respiratory period variability) not only strongly covaried across runs (Fig. 3A left panel), but also consistently exhibited excellent reliability, with ICCs between 0.75 and 0.94 (Fig. 3A right panel). Whether such a reliable noise-generating process also translates into a reliable influence on the measure of interest (i.e. gray matter time series data) was investigated next.

**Figure 3.**
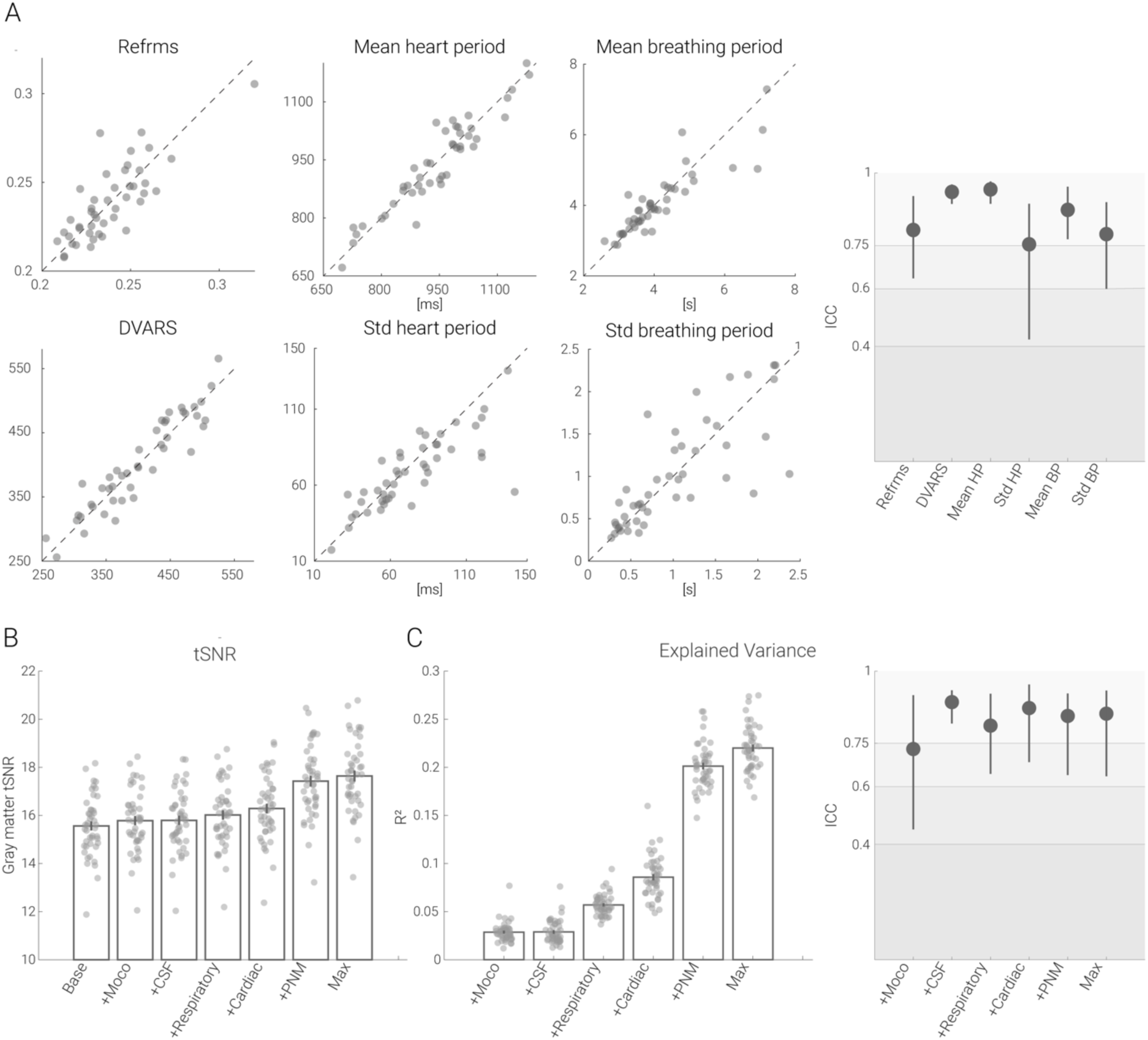
Reliability of physiological measurements and effects on tSNR and explained variance in the gray matter. **A.** Scatter plots show the metrics derived from physiological measurements recorded in each session, plotted against each other (session 1 on x-axis, session 2 on y axis) for every participant. On the very right, associated ICC values are depicted with the dots (lines depict 95% confidence intervals). **B.** Bar graphs show the gray matter tSNR after various physiological noise correction techniques have been applied. **C.** On the left, the bar graphs show the gray matter time series variance accounted for by various physiological noise correction techniques. In all bar plots, the vertical lines on the bars depict the standard error of the mean and the circles indicate participant-specific values. On the right, ICC values for explained variance are shown with the filled circles and the lines depicting 95% confidence intervals. The gray scale background reflects the ICC ranges (as defined by Cicchetti & Sparrow (1981) and Hallgren (2012)): poor <0.4, fair 0.4–0.59, good 0.6–0.74, excellent ≥0.75.

Therefore, we assessed the effects of noise sources on tSNR (an often-used metric of fMRI time series) and explained variance. With respect to gray matter tSNR changes (Figure 3B), the addition of the noise regressors led to the following increases: motion regressors 1.4%, CSF regressor 1.5%, respiratory regressors 2.9%, cardiac regressors 4.7%, PNM regressors 11.9%, and the combination of all regressors 13.4% (compared to the tSNR after the baseline pipeline), with all increases being significant at p < 0.001. Looking at this from the perspective of the fraction of gray matter time series variance explained by each of the noise regressors, we observed the following (Figure 3C right panel): motion regressors and the CSF regressor both 2.9%, respiratory and cardiac regressors 5.7% and 8.6%, PNM regressors 20.1% and combining all regressors 22.0%. Most importantly though, the variance explained by each of the noise components was highly reliable between runs (Figure 3C left panel): ICC values were mostly in the excellent range, varying between 0.73 to 0.89. Such a pattern of results is consistent with the above-mentioned reduction in amplitude and reliability of functional connectivity after denoising and provides evidence for the presence of structured and reliable non-neural signals being present in the gray matter time series.

#### 3.2.3. Thermal noise

After having assessed the impact of physiological noise, we now turn our focus to the influence of thermal noise. We aimed to remove thermal noise either via MP-PCA or via spatial smoothing – both of these approaches were added to the maximal denoising pipeline for physiological noise (more specifically, they occurred before GLM-based physiological denoising), which now also served as the baseline to compare against.

Since thermal noise removal has to our knowledge not been addressed in the spinal fMRI literature yet, we first assessed its impact on tSNR and observed a highly significant (all p < 0.001) increase in gray matter tSNR after adding either MP-PCA (140.2%) or spatial smoothing with a 2mm (120.2%, p < 0.001) or 4mm kernel (260.4%, p < 0.001). This increase in tSNR was thus similar to what was observed when adding physiological noise correction regressors, though now of much stronger amplitude. In sharp contrast to physiological noise correction however, both MP-PCA and spatial smoothing led to an increase in functional connectivity amplitudes (Table 3 and Figure 4): dorsal-dorsal, ventral-ventral and between-hemicord dorsal-ventral connectivity all had significantly higher amplitudes when compared to the maximal denoising pipeline; the absolute strength of within-hemicord dorsal-ventral connectivity also increased, though with a sign-change, which turned from negative to positive after MP-PCA and smoothing. For all connections, the reliability of functional connectivity increased when spatial smoothing was added to maximal denoising pipeline, whereas a more mixed picture appeared for MP-PCA (with either a slight decrease [dorsal-dorsal and ventral-ventral], increase [between-hemicord] or no change [within-hemicord]; Tables 3 and Figure 4).

**Figure 4.**
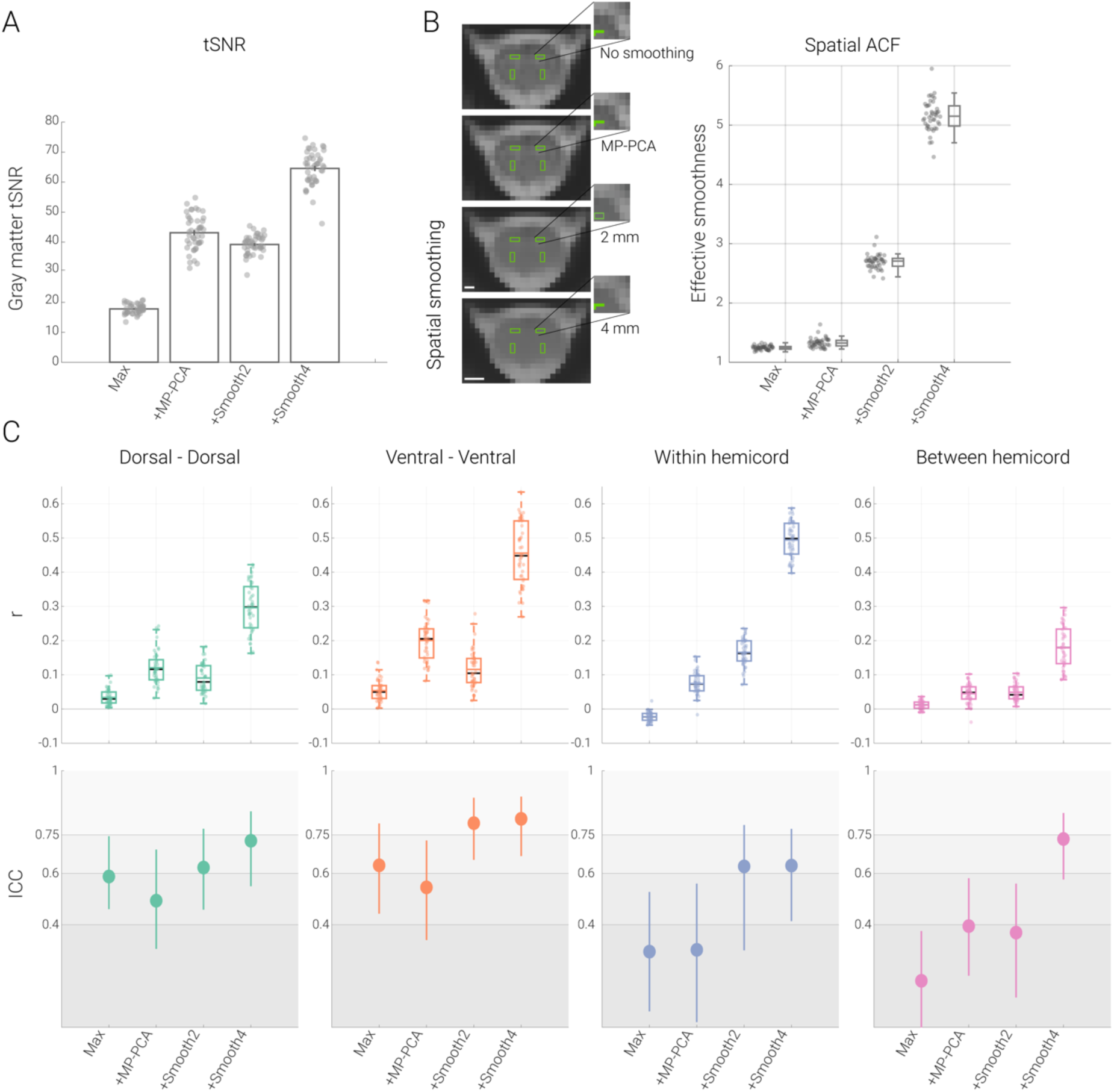
Impact of thermal noise removal. **A. Impact of thermal noise removal on tSNR.** Bar graph shows the tSNR in the gray matter for each segment after employing different processing pipelines (Max: maximal processing – which served as baseline for this comparison, +MP-PCA: maximal + thermal noise removal via MP-PCA; +Smooth2: maximal + smoothing with a 2mm kernel; +Smooth4: maximal + smoothing with a 4mm kernel). The vertical lines on the bars depict the standard error of the mean and the filled dots lines indicate participant-specific values. **B. Impact of thermal noise removal on spatial smoothness.** On the left side, one exemplary EPI slice of a participant in native space (where analyses were carried out) and gray matter ROIs overlaid in green are shown after different processing steps. Scale bars represent 2mm and 4mm, respectively. On the right side, effective spatial smoothness values estimated using AFNI’s 3dFWHMx function are depicted via box-plots for which the median is denoted by the central mark and the bottom and top edges of the boxes represent the 25th and 75th percentiles, respectively, with the whiskers encompassing ∼99% of the data. The circles represent individual participants. **C. Impact of thermal noise removal on functional connectivity and reliability.** The top panel depicts Pearson correlation values (averaged across two sessions) between the time-courses of different ROIs with the box plots for four different pipelines (box plots are identical to those in B – except here the mean is denoted by the colored central mark). On the bottom panel, ICC values for each connection (and each pipeline) are shown with the filled circles and the lines show 95% confidence intervals. The gray scale background reflects the ICC ranges (as defined by Cicchetti & Sparrow (1981) and Hallgren (2012)): poor <0.4, fair 0.4–0.59, good 0.6–0.74, excellent ≥0.75.

**Table 3.** Functional connectivity and its reliability after thermal noise correction procedures. This table depicts functional connectivity and reliability results of each connection for different thermal noise correction processing pipelines. r represents the mean Pearson correlation across participants, t and p represent the t-value and two-tailed FWE-corrected (for four tests) p-value from a permutation test (against 0), respectively. ICC(95% CI) represents ICC(2,1) values and 95% bootstrapped confidence intervals.

|  | **Dorsal**<br>**Dorsal** | **Ventral**<br>**Ventral** | **Within**<br>**Hemicord** | **Between**<br>**Hemicord** |
| --- | --- | --- | --- | --- |
| **Maximal** | r = 0.03<br>t = 9.5<br>p < 0.001<br>ICC (95% CI) =<br>0.59 (0.46 – 0.74) | r = 0.05<br>t = 11.6<br>p < 0.001<br>ICC (95% CI) =<br>0.63 (0.44 – 0.79) | r = -0.02<br>t = -10.7<br>p < 0.001<br>ICC (95% CI) =<br>0.30 (0.06 – 0.53) | r = 0.01<br>t = 6.7<br>p < 0.001<br>ICC (95% CI) =<br>0.18 (-0.03 – 0.38) |
| **Thermal noise**<br>**removal +**<br>**maximal** | r = 0.12<br>t = 16.7<br>p < 0.001<br>ICC (95% CI) =<br>0.49 (0.31 – 0.69) | r = 0.20<br>t = 22.9<br>p < 0.001<br>ICC (95% CI) =<br>0.55 (0.34 – 0.73) | r = 0.07<br>t = 15.6<br>p < 0.001<br>ICC (95% CI) =<br>0.30 (0.02 – 0.56) | r = 0.05<br>t = 10.6<br>p < 0.001<br>ICC (95% CI) =<br>0.39 (0.20 – 0.58) |
| **Maximal +**<br>**2mm**<br>**smoothing** | r = 0.09<br>t = 14.3<br>p < 0.001<br>ICC (95% CI) =<br>0.62 (0.46 – 0.77) | r = 0.12<br>t = 13.9<br>p < 0.001<br>ICC (95% CI) =<br>0.79 (0.65 – 0.89) | r = 0.17<br>t = 29.1<br>p < 0.001<br>ICC (95% CI) =<br>0.63 (0.30 – 0.79) | r = 0.05<br>t = 13.4<br>p < 0.001<br>ICC (95% CI) =<br>0.37 (0.12 – 0.56) |
| **Maximal +**<br>**4mm**<br>**smoothing** | r = 0.29<br>t = 28.3<br>p < 0.001<br>ICC (95% CI) =<br>0.73 (0.55 – 0.84) | r = 0.46<br>t = 33.6<br>p < 0.001<br>ICC (95% CI) =<br>0.81 (0.67 – 0.89) | r = 0.49<br>t = 64.4<br>p < 0.001<br>ICC (95% CI) =<br>0.63 (0.41 – 0.77) | r = 0.18<br>t = 19.8<br>p < 0.001<br>ICC (95% CI) =<br>0.73 (0.58 – 0.84) |

One aspect of these results deserves further interrogation, namely whether the increased connectivity amplitudes might simply come about via time-course mixing between the ROIs due to an increased spatial smoothness of the data after the thermal noise correction procedures. We therefore assessed the spatial autocorrelation function of the EPI data and observed that – across the group – the effective smoothness increased from 1.2±0.03 by 116% for 2mm (2.7±0.12) and 313% for 4mm (5.2±0.26) smoothing. Importantly, despite the more than two-fold increase in tSNR and connectivity amplitudes observed after MP-PCA, this procedure only led to a 7% increase in spatial smoothness (1.3±0.08). It is thus unlikely that the increased connectivity observed after MP-PCA is driven via time-course mixing between the different ROI – an assumption underscored even further by the fact the MP-PCA increased the connectivity of all connections in a way that is unrelated to the ROIs’ spatial distance (Figure S3). Conversely, the effects of spatial smoothing on connectivity amplitudes are likely driven by time-course mixing, since i) the largest increase e.g. for 2mm smoothing was observed for the ROIs being closest together (dorsal-ventral within-hemicord connection; Figure S3) and ii) the increase in connectivity parallels the increase in spatial smoothness (cf. Figure 4B and 4C). This suggests that even modest smoothing kernels such as 2mm should only be employed with great caution in the spinal cord.

### 3.3. Within-segment functional connectivity

Finally, we aimed to assess whether resting-state functional connectivity could also be reliably observed at the level of single spinal segments (C3, C4, C5, C6, C7, C8 and T1; Figure 5A). For these analyses we used data that were denoised with MP-PCA in addition to the maximal physiological noise correction pipeline, as the above analyses showed this method to be beneficial for both tSNR and connectivity estimates.

**Figure 5.**
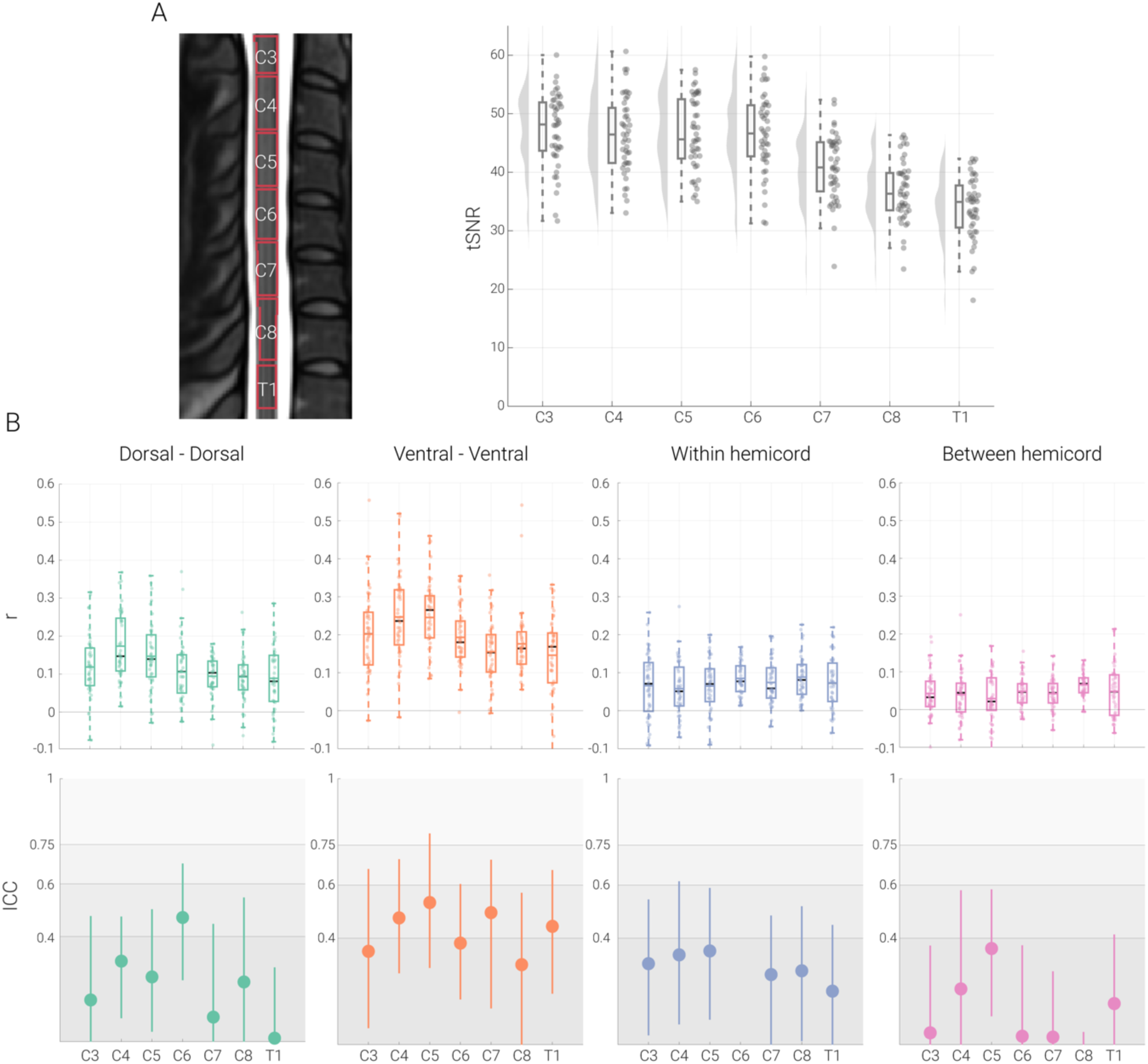
Segment-specific functional connectivity. **A.** The midsagittal cross-section on the left (from the T2-weighted PAM50 template image) shows the thresholded probabilistic segments overlaid as outlines. Segment-wise tSNR values are depicted via box-plots for which the median is denoted by the central mark and the bottom and top edges of the boxes represent the 25th and 75th percentiles, respectively, with the whiskers encompassing ∼99% of the data. The circles represent individual participants and half-violin plots show the distribution across participants. **B.** The top panel depicts Pearson correlation values (averaged across two sessions) between different ROIs with one box plot per segmental level. For the box plots, the median and mean are denoted by the central black mark and the colored mark, respectively. The bottom and top edges of the boxes represent the 25th and 75th percentiles, respectively, with the whiskers encompassing ∼99% of the data. The circles represent individual participants. The bottom panel depicts ICC values for each connection with the dot and the lines denote 95% confidence intervals; please note that the ICC for within hemicord connectivity for level C6 is far below zero, resulting in it not being visible here (see Table 4 for all ICC values). The gray scale background reflects the ICC ranges (as defined by Cicchetti & Sparrow (1981) and Hallgren (2012)): poor <0.4, fair 0.4–0.59, good 0.6–0.74, excellent ≥0.75.

**Table 4.** Functional connectivity and its reliability for different spinal segments. This table depicts functional connectivity and reliability results of each connection at different spinal segments. r represents the mean Pearson correlation across participants, t and p represent the t-value and two-tailed family-wise-error corrected p-value from a permutation test (against 0), respectively. ICC (95% CI) represents ICC(2,1) values and 95% bootstrapped confidence intervals.

|  | **Dorsal**<br>**Dorsal** | **Ventral**<br>**Ventral** | **Within**<br>**Hemicord** | **Between**<br>**Hemicord** |
| --- | --- | --- | --- | --- |
| **C3** | r = 0.12<br>t = 9.2<br>p < 0.001<br>ICC (95% CI) =<br>0.16 (-0.24 – 0.48) | r = 0.20<br>t = 8.9<br>p < 0.001<br>ICC (95% CI) =<br>0.35 (0.06 – 0.66) | r = 0.07<br>t = 5.0<br>p < 0.001<br>ICC (95% CI) =<br>0.30 (0.03 – 0.55) | r = 0.04<br>t = 4.6<br>p < 0.001<br>ICC (95% CI) = 0.04<br>(-0.23 – 0.37) |
| **C4** | r = 0.17<br>t = 13.6<br>p < 0.001<br>ICC (95% CI) =<br>0.31 (0.09 – 0.48) | r = 0.25<br>t = 13.3<br>p < 0.001<br>ICC (95% CI) =<br>0.48 (0.27 – 0.69) | r = 0.06<br>t = 5.5<br>p < 0.001<br>ICC (95% CI) =<br>0.34 (0.08 – 0.61) | r = 0.04<br>t = 4.2<br>p < 0.001<br>ICC (95% CI) =<br>0.21 (-0.07 – 0.58) |
| **C5** | r = 0.15<br>t = 10.9<br>p < 0.001<br>ICC (95% CI) =<br>0.25 (0.04 – 0.50) | r = 0.25<br>t = 12.5<br>p < 0.001<br>ICC (95% CI) =<br>0.53 (0.29 – 0.79) | r = 0.06<br>t = 5.7<br>p < 0.001<br>ICC (95% CI) =<br>0.35 (0.09 – 0.59) | r = 0.03<br>t = 2.6<br>p = 0.07<br>ICC (95% CI) = 0.36<br>(0.11 – 0.58) |
| **C6** | r = 0.11<br>t = 7.5<br>p < 0.001<br>ICC (95% CI) =<br>0.47 (0.23 – 0.68) | r = 0.19<br>t = 16.3<br>p < 0.001<br>ICC (95% CI) =<br>0.38 (0.17 – 0.60) | r = 0.09<br>t = 12.4<br>p < 0.001<br>ICC (95% CI) = -<br>0.24 (-0.52 – 0.0) | r = 0.05<br>t = 7.5<br>p < 0.001<br>ICC (95% CI) = 0.03<br>(-0.45 – 0.37) |
| **C7** | r = 0.09<br>t = 11.5<br>p < 0.001<br>ICC (95% CI) =<br>0.09 (-0.29 – 0.45) | r = 0.16<br>t = 18.2<br>p < 0.001<br>ICC (95% CI) =<br>0.49 (0.14 – 0.69) | r = 0.07<br>t = 8.1<br>p < 0.001<br>ICC (95% CI) =<br>0.26 (0.0 – 0.49) | r = 0.04<br>t = 7.2<br>p < 0.001<br>ICC (95% CI) = 0.03<br>(-0.23 – 0.28) |
| **C8** | r = 0.049<br>t = 9.5<br>p < 0.001<br>ICC (95% CI) =<br>0.23 (-0.20 – 0.55) | r = 0.18<br>t = 16.6<br>p < 0.001<br>ICC (95% CI) =<br>0.30 (-0.13 – 0.57) | r = 0.09<br>t = 9.9<br>p < 0.001<br>ICC (95% CI) =<br>0.28 (0.01 – 0.52) | r = 0.07<br>t = 14.0<br>p < 0.001<br>ICC (95% CI) = -<br>0.23 (-0.46 – 0.04) |
| **T1** | r = 0.09<br>t = 7.0<br>p < 0.001<br>ICC (95% CI) =<br>0.01 (-0.22 – 0.28) | r = 0.15<br>t = 12.2<br>p < 0.001<br>ICC (95% CI) =<br>0.44 (0.19 – 0.66) | r = 0.07<br>t = 6.9<br>p < 0.001<br>ICC (95% CI) =<br>0.20 (-0.09 – 0.45) | r = 0.05<br>t = 4.8<br>p < 0.001<br>ICC (95% CI) = 0.15<br>(-0.09 – 0.41) |

First of all, we observed that – despite the use of z-shimming – the gray matter tSNR was lower for the lowermost segments (C7, C8 and T1). Functional connectivity, however, was highly significant in every segment for all connections (dorsal-dorsal, ventral-ventral, within-hemicord, between-hemicord; see Figure 5 and Table 4). Reliability of functional connectivity at the single-segment level, on the other hand, was mostly poor (see Figure 5 and Table 4). For dorsal-dorsal connectivity, the reliability values were largely in the poor range except at level C6 (in the fair range), and for ventral-ventral connectivity, the ICC values fluctuated between the poor and fair range (poor for C3, C6 and C8; fair for C4, C5, C7 and T1). Within- and between-hemicord dorsal-ventral reliability values were in the poor range for every single segment. These results highlight that even though it is possible to detect single-segment connectivity patterns, these are highly variable across scan-sessions and thus lack robustness with the currently employed approaches for data acquisition and analysis.

## 4. Discussion

In the last decade, evidence has accumulated that the human spinal cord exhibits spatially distinct patterns of spontaneous activity at rest, as functional connectivity was observed to exist between the two dorsal horns and between the two ventral horns, mirroring the functional division of the gray matter into sensory and motor parts, respectively. While this has generated interest in the use of such connectivity metrics in the clinical context as possible biomarkers for sensory and motor disorders (such as chronic pain and multiple sclerosis), a first essential step is to quantify their reliability, which we set out to do here at the clinically relevant field strength of 3T. We first replicated and extended previous resting-state fMRI findings by investigating the spinal cord’s functional connectivity and assessing its test-retest reliability in a large sample (N > 40). Considering that spinal cord BOLD signals are strongly affected by noise, we characterized the impact of various noise sources (i.e., physiological noise and thermal noise) on connectivity strength and reliability. Finally, we considered local aspects of functional connectivity and their reliability by investigating this at a macro-scale unit of spinal cord organization, namely at the level of single spinal segments.

### 4.1. Replication and extension of previous resting-state functional connectivity results

In order to replicate previously observed functional connectivity results, we used a commonly employed processing pipeline for removal of physiological noise (i.e. addressing noise arising from participant motion, cardiac, respiratory and CSF effects). With an ROI-based approach, we demonstrated statistically significant functional connectivity between the dorsal horns (housing somatosensory function) and between the ventral horns (housing somatomotor function), thus replicating a pattern of results observed in previous spinal cord fMRI studies in rats (Wu et al., 2018), monkeys (Chen et al., 2015; Wu et al., 2019) and humans (3T: Barry et al., 2018; Eippert et al., 2017b; Hu et al., 2018; Liu et al., 2016; Weber et al., 2018 ; 7T: Barry et al., 2014, 2016; Conrad et al., 2018). The fact that such a functional connectivity profile is observed across different acquisition protocols, field strengths as well as species provides further support for the hypothesis that intrinsic fluctuations of the spinal cord are not of random nature. It does however neither confirm the neuronal origin of resting-state functional connectivity nor provide answers regarding the exact neurobiological underpinnings (Eippert & Tracey, 2014) and towards this end, combining fMRI with electrophysiological recordings (Brookes et al., 2011; Schölvinck et al., 2010) would be beneficial, with important first steps in this direction already being taken (Wu et al., 2019).

We also observed significant functional connectivity within (left dorsal-ventral and right dorsal-ventral) and between (left dorsal - right ventral and right dorsal - left ventral) hemicords, though these were clearly weaker in terms of correlation magnitude than the dorsal-dorsal and ventral-ventral connections (and were actually negative for within-hemicord connectivity). This weaker result observed here fits well into the literature, with some studies observing similar sensory-motor cord connectivity (Chen et al., 2015; Weber et al., 2018; Wu et al., 2019), and others not (Barry et al., 2014; Eippert et al., 2017a; see Harrison et al., 2021 for a review). Of note in this case are recent electrophysiological data providing evidence for such dorsal-ventral connectivity at the level of local field potentials and spike trains in anaesthetized animals (McPherson & Bandres, 2021; Wu et al., 2019). While the reason for this variability of *functional* connectivity findings across experimental models and measurement-levels is currently unclear, existence for *structural* dorsal-ventral connectivity is unequivocal, as it is the anatomical substrate for polysynaptic spinal reflexes in humans (Pierrot-Deseilligny & Burke, 2012; Sandrini et al., 2005) and has also been delineated in detail with modern tracing approaches in mice (e.g. Ronzano et al., 2021; Stepien et al., 2010). Interestingly, in the context of fMRI, the likelihood to observe dorsal-ventral resting-state connectivity might also depend on data processing choices, as this type of result is not robust against variations in the processing pipeline (Eippert et al., 2017a; similar to what we observed here after removal of thermal noise).

One further way to judge the robustness of results is via their reliability, which we assessed here via test-retest reliability (Shrout & Fleiss, 1979). Using ICC as a measure of reliability, we observed fair-to-good reliability for dorsal-dorsal and ventral-ventral connectivity and poor reliability for within hemicord and between hemicord connectivity (the robustness of this finding received further support from analyses in which we employed partial correlation instead of Pearson correlation and observed highly similar results). This is in line with a previous investigation by Barry and colleagues (2016) at the ultra-high field strength of 7T and demonstrates that a similar level of reliability can be obtained at the clinically-relevant field strength of 3T. Previous important investigations into the test-retest reliability of functional connectivity at 3T were limited in terms of the employed sample size (N=10 for Liu et al. (2016), Hu et al. (2018), Barry et al. (2018)), which we overcame here using a more than 4-fold larger sample size. Other studies have assessed the split-half reliability of ICA-derived spinal cord resting-state networks in humans at 3T (Kong et al., 2014) and the test-retest reliability of ROI-based functional connectivity in rats at 9.4T (Wu et al., 2018) and generally observed fair to good reliability as well. It is important to point out that despite these differences in data acquisition and analyses – which have been demonstrated to substantially influence reliability estimates of resting-state connectivity in the brain (for review, see Noble et al. (2019)) – all of these findings seem to point towards reproducible results, i.e. show the presence of reliable spinal cord resting-state networks.

### 4.2. Impact of noise sources on resting-state functional connectivity and its reliability

Considering that noise has an immense impact on the spinal cord fMRI signal – i.e. its influence is much more prominent than in the brain (Piche et al., 2009; Cohen-Adad et al., 2010) – we next assessed to what degree functional connectivity and its reliability are affected by various noise sources and procedures for their correction.

We first investigated the impact of physiological noise regression on functional connectivity and observed that, in general, extensive denoising (i.e. the addition of various physiological noise regressors to the baseline) led to a clear decrease in the amplitude of functional connectivity estimates and also decreased the reliability of functional connectivity, while – not surprisingly – tSNR was increased. This reduction in amplitude and reliability may seem counterintuitive at first glance, as one might expect that removal of physiological noise should improve the detectability and reliability of functional connectivity. However, this result is indeed consistent with observations in many resting-state fMRI studies in the brain (Birn et al., 2014; Guo et al., 2012; Noble et al., 2019; Parkes et al., 2018; Shirer et al., 2015; Zou et al., 2015), where a decrease in reliability was observed after various denoising approaches.

Further investigations undertaken to elucidate why reliability decreased after physiological noise removal revealed that the sources of physiological noise – e.g. mean and standard deviation of heart period and breathing period – were highly reliable, i.e. showed stable responses within participants across runs, but large variation across participants (in this sense, we are removing ‘true’ biological variability here, though of a confounding nature). The same held for the amount gray matter time series variance explained by physiological noise regressors: these mostly exhibited reliability in the excellent range, in line with observations in previous studies that also looked at the reproducibility of respiratory and cardiac effects in spinal cord MRI data (Piché et al., 2009; Verma & Cohen-Adad, 2014). If one now considers that our reliability metric of choice – the ICC – can be roughly defined as a ratio of the variance of interest (in our case: between-participant) to the total variance (Liljequist et al., 2019), a possible path via which physiological noise removal decreases reliability becomes apparent: it removes spatiotemporally structured ‘reliable artefacts’ (i.e. differing strongly between participants, but not necessarily between runs within participants), that would otherwise contribute to the reliability estimation via their confounding effects on connectivity. A similar argument has already been made for the reliability of resting-state connectivity in the brain, substantiated by a detailed investigation of the changes in the different variance components contributing to the ICC (Birn et al., 2014). In other words: once the impact of these *reliable* non-neural sources that influence ROI time-courses similarly – and thus also increase the correlation strength – within each participant is removed, correlation amplitude as well as reliability decreases.

Thus, and as already pointed out by others (Birn et al., 2014; Shirer et al., 2015; Noble et al., 2019), the reduction in reliability after physiological noise removal might actually increase the validity of the results. Validity can be defined as how close or accurate one is measuring what one intends to measure (Carmines & Zeller, 1979) and in our case – using resting-state fMRI – we intend to measure neuronally driven BOLD fluctuations, which only represent a small percentage of the variance in the noisy fMRI signal (Bijsterbosch & Beckmann, 2017; Birn, 2012). One might anticipate that an improved validity after removal of physiological noise may also lead to a better distinction at the group level – e.g. between patients’ and healthy controls’ functional connectivity patterns – or improve the relationship between functional connectivity estimates and ‘trait’ characteristics (Shirer et al., 2015; Noble et al., 2017a; Noble et al., 2019); interventional studies could also shed light on this.

In addition to the effects of removing physiological noise, we also assessed the impact of thermal noise (Edelstein et al., 1986; Hoult & Richards, 1976; Krüger & Glover, 2001) and methods for its correction. While we did not formally assess the physiological noise to thermal noise ratio in our data – as this depends on many factors (Brooks et al., 2013; Triantafyllou et al., 2005, 2011) and is complicated by the fact that part of what is traditionally considered physiological ‘noise’ is our signal of interest here – we observed marked effects of thermal noise removal: the application of MP-PCA (Veraart et al., 2016a; Veraart et al., 2016b) led to i) a substantial increase in tSNR (more than two-fold), ii) a concurrent and consistent increase in correlation strength (more than three-fold) and iii) no consistent changes in reliability (as we observed either decreases, no change or an increase in reliability, possibly warranting future investigations). One immediately notices the clear difference to physiological noise removal, which also increased the tSNR, but decreased connectivity strength and reliability, likely due to physiological noise being structured and reliable. Despite being a major source of noise in fMRI acquisitions, only a few brain fMRI studies (Ades-Aron et al., 2021a; Adhikari et al., 2019) utilized thermal noise removal via MP-PCA and to our knowledge its benefits for spinal cord fMRI had not yet been demonstrated (see Grussu et al., 2020 for an application of MP-PCA in quantitative MRI of the cord and Vizioli et al., 2021 for an even more recent thermal noise correction technique applied to brain fMRI data). We furthermore compared MP-PCA to spatial smoothing which also serves to suppress thermal noise: compared to spatial smoothing (which also enhanced tSNR and connectivity strength), MP-PCA achieved this without incurring a substantial penalty in terms of increased spatial smoothness. This is an important consideration, since ROIs in the spinal cord lie so close to each other that even with a modest Gaussian smoothing kernel of 2mm FWHM, artificial connectivity (via time-course mixing) can be induced, which we were able to demonstrate here, since the increase in connectivity strength induced via smoothing depended on the spatial proximity of the ROIs. We thus believe that thermal noise removal via MP-PCA might be an attractive option for enhancing the sensitivity of spinal cord fMRI, but would like to note that its detailed validation in the context of resting-state fMRI is still outstanding (as are comparisons with other methods, e.g. Vizioli et al., 2021; Dowdle et al., 2023).

### 4.3. Within segment functional connectivity

Finally, we assessed the amplitude and reliability of more localized aspects of connectivity, i.e. within a spinal cord segment, which is traditionally considered to be the basic organizational unit of the spinal cord along the rostrocaudal axis (though see Watson & Sidhu, 2009; Sengul et al., 2013). This was made possible by the availability of probabilistic maps for spinal cord segments (Cadotte et al., 2015) and their integration into a common template space (De Leener et al., 2017). Reassuringly, for all of the segmental levels that we investigated (C3-T1), we were able to demonstrate robust functional connectivity patterns, i.e. significantly positive correlations between bilateral dorsal and between bilateral ventral horns, despite an apparent decrease in tSNR for segments C7-T1 compared to the more rostral cervical segments. While minor variations in connectivity strength were observed, the overall pattern stayed consistent across segments and mirrored the above-reported connectivity results that spanned the superior-inferior axis of the imaging volume (similar to Eippert et al. (2017a)). We also observed significant within and between hemicord dorsal-ventral connectivity at each segment (except C5 where between hemicord connectivity was not significant), though this was again much weaker than dorsal-dorsal and ventral-ventral connectivity. Importantly though, the reliability of functional connectivity at the level of individual segments was consistently in the poor range: this held entirely for dorsal-ventral connectivity, mostly for dorsal-dorsal connectivity (apart from segment C6) and partially for ventral-ventral connectivity (where approximately half of the ICCs were in the fair range); in addition, this was consistently evident across segments and thus not driven by the lower tSNR present in the more caudal segments. Given our 5mm slice thickness, there were only approximately three EPI slices in each segment, probably rendering correlation estimates susceptible to remaining noise across voxels (e.g. compared to the analyses across the imaging volume) and recent investigations have suggested that other 3T acquisition approaches might be helpful in this regard (Kinany et al., 2022), as could be the use of higher field strength (Barry et al., 2018) or using slightly dilated regions of interest. Considering that many disorders present with localized spinal cord pathology (e.g. cervical myelopathy; Nouri et al., 2015) and that spinal cord resting-state fMRI is now being applied in such contexts – e.g. spinal cord injury (Chen et al., 2015; Sengupta et al., 2021) or multiple sclerosis (Conrad et al., 2018; Combes et al., 2022) – it will be of utmost importance to improve the reliability of segment-wise connectivity via optimization of data acquisition and analysis approaches, since only with a reliable estimate of connectivity can longitudinal studies that monitor disease progression or treatment effects be carried out successfully.

### 4.4. Limitations and outlook

There are several limitations of the current study that are worth mentioning. *First* of all, in terms of assessing functional connectivity, we have only used ROI-based static functional connectivity approaches here, whereas data-driven approaches like ICA (Kong et al., 2014) or time-varying functional connectivity approaches (Kinany et al., 2020) might yield different insights into the reliability of spinal cord networks; of note, these could be applied on our openly-available data-set, allowing for a direct comparison between methods. *Second*, we assessed the impact of physiological noise solely within the PNM framework (Brooks et al., 2008; Kong et al., 2012). Although PNM is well established for spinal cord fMRI and has compared favorably against other methods in this context (Kong et al., 2012), there are many other approaches to address physiological noise that we did not consider here and that again might perform differently, such as CompCor (Behzadi et al., 2007), DRIFTER (Särkkä et al., 2012) or ICA-AROMA (Pruim et al., 2015). A comparison of various denoising approaches was beyond the scope of current work (similar to evaluating the effects of different preprocessing steps), but could also be carried out on this openly-available data-set and might offer additional insights, as there might be unmodeled noise components still present in the data. *Third*, considering the various different approaches for data acquisition that are currently employed in spinal cord fMRI at 3T (e.g. Barry et al., 2021; Kinany et al., 2022), we refrain from extrapolating our results beyond the specific acquisition scheme employed here. *Fourth*, one needs to be careful regarding the interpretation of the observed reliability, since on the one hand, our results may represent an ‘upper’ end of reliability estimates, as we assessed the test-retest reliability of functional runs which were separated by at most ∼10 minutes (see Kowalczyk et al., 2023 for an assessment of reliability between sessions). On the other hand, the two functional runs had slightly different z-shim settings which might bias towards ‘lower’ reliability (although there were no significant tSNR differences between the two acquisitions). Given these factors, it would be interesting to assess the reliability of resting-state spinal networks over different time spans in the future, ranging from hours to days to months, as reliability may decrease over time (Shehzad et al., 2009) – here one could also envision to assess sessions that were acquired in different scanners (Noble, et al., 2017b) in order to probe different components of reliability (Brandmaier et al., 2018). *Fifth*, all connectivity results reported here are based on within-slice correlations, i.e. we did not address rostro-caudal time series correlations and an assessment of the full correlation matrix (including between-segment connectivity) remains for future studies. *Finally*, it is important to keep in mind that the ICC is calculated as a ratio of between person variance to total variance and ICC values are thus dependent on the characteristics of given sample. For instance, ICC values for patient groups (such as multiple sclerosis or chronic pain) might be higher due to the larger variability between individual patients as compared to our very homogenous sample consisting of young healthy adults in a very restricted age-range (see also Wenger et al., 2022). Consideration of these aspects might be helpful for understanding the limitations and benefits of spinal cord resting-state fMRI in the clinical context where longitudinal as well as multi-site and multi-cohort studies are common.

## 5. Conclusion

Taken together, this study adds to a growing body of evidence that the spinal cord exhibits structured resting-state functional connectivity. Connectivity within sensory and within motor regions of the spinal cord seems to be of robust nature, as it presents with fair-to-good reliability. Our results furthermore underscore the critical need for addressing physiological noise, though now from the perspective of reliability and also demonstrate that thermal noise removal can have beneficial effects on the detection of functional connectivity. Finally, our assessments of segment-level connectivity (presenting with low reliability) provide a more cautionary note and suggest that further improvements in data acquisition and analysis would be important before employing resting-state spinal cord fMRI longitudinally in the context of assessing disease progression or treatment response.

## Acknowledgments

JB received funding from the UK Medical Research Council (MR/N026969/1). FE received funding from the Max Planck Society and the European Research Council (under the European Union’s Horizon 2020 research and innovation programme; grant agreement No 758974). NW received funding from the European Research Council under the European Union’s Seventh Framework Programme (FP7/2007-2013, ERC grant agreement No 616905), the European Union’s Horizon 2020 research and innovation programme (under the grant agreement No 681094) and the BMBF (01EW1711A & B) in the framework of ERA-NET NEURON. The authors would like to thank Alice Dabbagh for helpful discussions on fMRI reliability, the radiographers at MPI CBS for invaluable help with data acquisition and all volunteers for taking part in this study, as well as Benjamin Ades-Aron and Paul Taylor for help with MP-PCA and 3dFWHMx implementation, respectively.

## Supplementary Material

**Figure S1.**
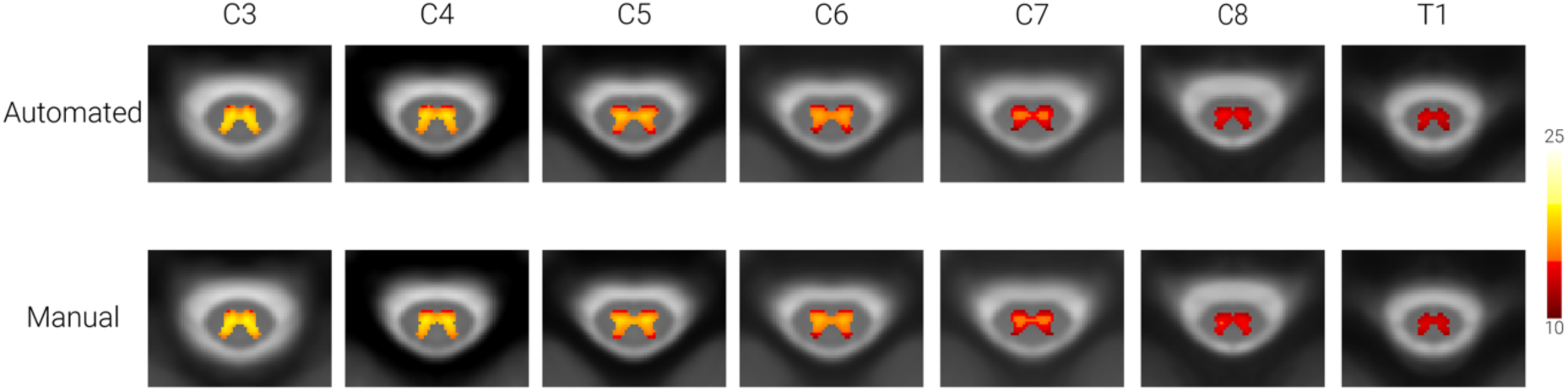
tSNR maps. Group-average gray matter tSNR maps of transversal slices at the middle of each segment are shown for the two different shim conditions (top row: Automated; bottom row: Manual). The maps are overlaid onto the PAM50 T2*-weighted template and depict a tSNR range from 10-25.

**Figure S2.**
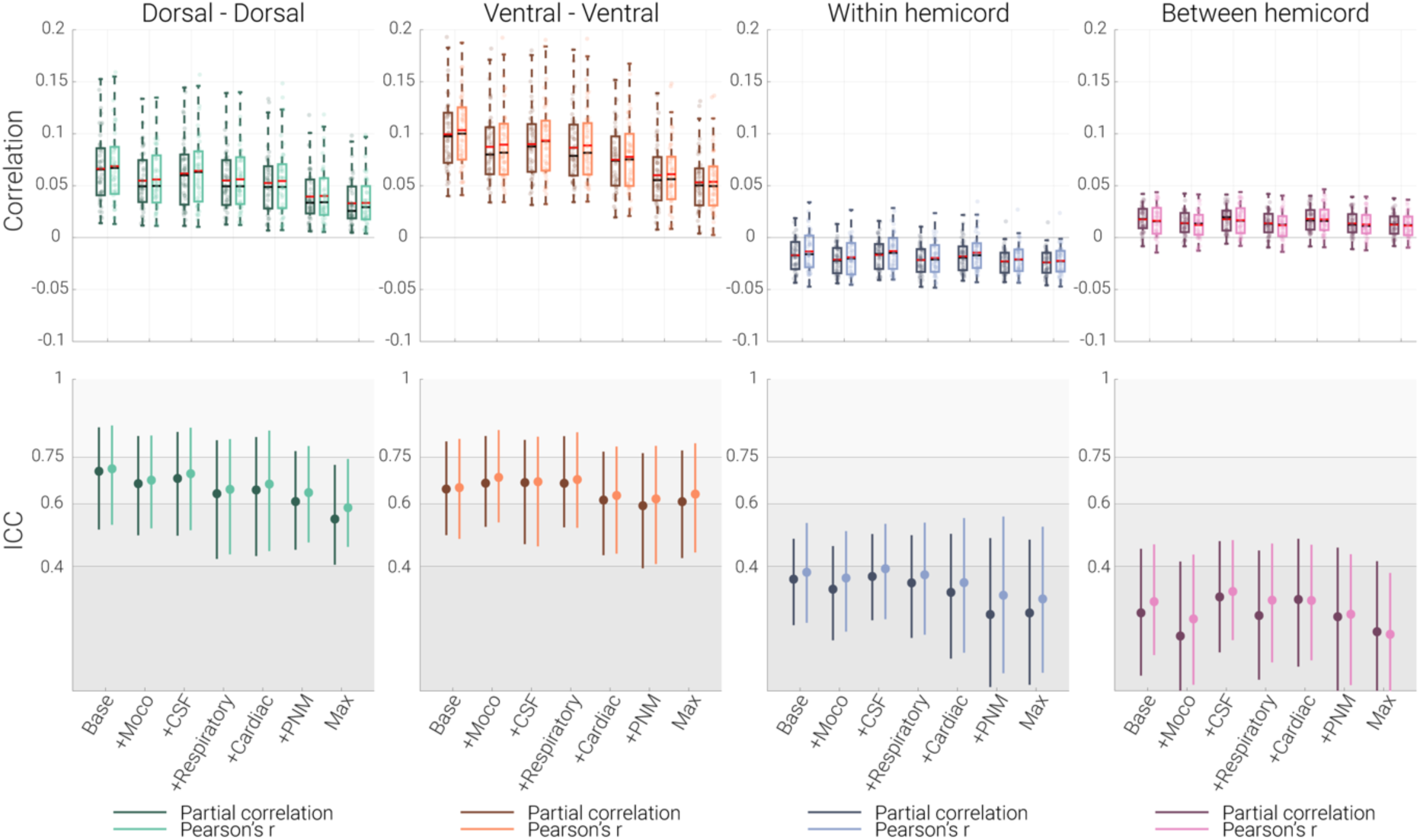
Partial correlation vs Pearson correlation. The top panels depict functional connectivity estimates between different ROIs calculated with either partial correlation or Pearson correlation (average across two sessions) using grouped box plots for the seven denoising pipelines. For the box plots, median and mean are denoted by the central black and red marks, respectively. The bottom and top edges of the boxes represent the 25th and 75th percentiles, respectively, with the whiskers encompassing ∼99% of the data. The bottom panels depict ICC values for different denoising pipelines with dots and lines denote 95% confidence intervals. The gray scale background reflects the ICC ranges (as defined by Cicchetti & Sparrow (1981) and Hallgren (2012)): poor <0.4, fair 0.4–0.59, good 0.6–0.74, excellent ≥0.75.

**Figure S3.**
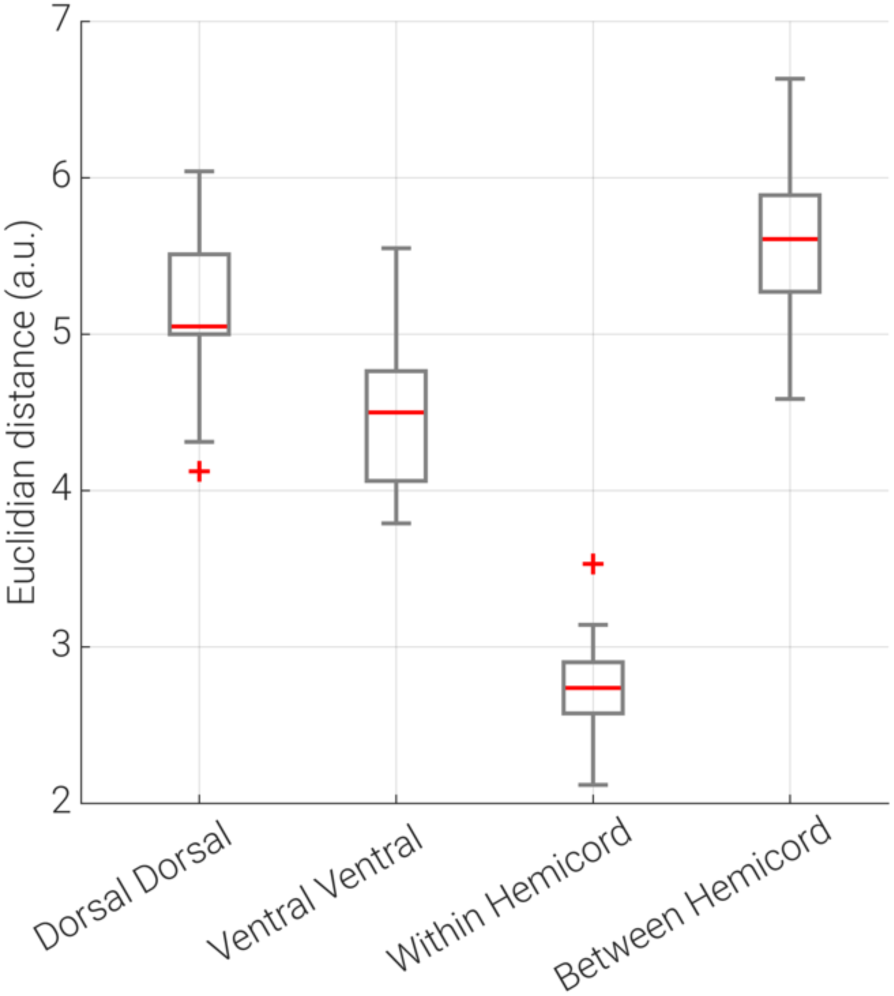
Euclidian distance between ROIs. Box plots show the median Euclidian distance between the closest voxels of different ROIs (within each slice) across slices and participants. The median is denoted by the central red line. The bottom and top edges of the boxes represent the 25th and 75th percentiles, respectively, with the whiskers encompassing ∼99% of the data, and the outliers are denoted with the red crosses.

**Table S1.** Functional connectivity and its reliability after addition of pre-whitening to the maximal denoising pipeline. This table depicts functional connectivity and reliability results of each connection for two processing pipelines: the maximal pipeline and the maximal pipeline with the inclusion of FILM pre-whitening. r represents the mean Pearson correlation across participants, t and p represent the t-value and two-tailed family-wise-error corrected p-value from a permutation test (against 0), respectively. ICC (95% CI) represents ICC(2,1) values and 95% bootstrapped confidence intervals.

|  | **Dorsal**<br>**Dorsal** | **Ventral**<br>**Ventral** | **Within**<br>**Hemicord** | **Between**<br>**Hemicord** |
| --- | --- | --- | --- | --- |
| **Maximal** | $r = 0.03$<br>$t = 9.5$<br>$p < 0.001$<br>ICC (95% CI) =<br>0.59 (0.46 – 0.74) | $r = 0.05$<br>$t = 11.6$<br>$p < 0.001$<br>ICC (95% CI) =<br>0.63 (0.44 – 0.79) | $r = -0.02$<br>$t = -10.7$<br>$p < 0.001$<br>ICC (95% CI) =<br>0.30 (0.06 – 0.53) | $r = 0.01$<br>$t = 6.7$<br>$p < 0.001$<br>ICC (95% CI) =<br>0.18 (-0.03 – 0.38) |
| **Maximal +**<br>**Pre-whitening** | $r = 0.03$<br>$t = 9.6$<br>$p < 0.001$<br>ICC (95% CI) =<br>0.59 (0.46 – 0.74) | $r = 0.05$<br>$t = 11.7$<br>$p < 0.001$<br>ICC (95% CI) =<br>0.65 (0.47 – 0.80) | $r = -0.02$<br>$t = -10.6$<br>$p < 0.001$<br>ICC (95% CI) =<br>0.29 (0.06 – 0.52) | $r = 0.01$<br>$t = 6.7$<br>$p < 0.001$<br>ICC (95% CI) =<br>0.17 (-0.05 – 0.37) |

**Table S2.** Functional connectivity and its reliability after addition of white matter regression to the baseline and maximal denoising pipelines. This table depicts functional connectivity and reliability results of each connection for four processing pipelines: the baseline pipeline, the baseline pipeline with the inclusion of white matter regression, the maximal pipeline and the maximal pipeline with the inclusion of white matter regression. r represents the mean Pearson correlation across participants, t and p represent the t-value and two-tailed family-wise-error corrected p-value from a permutation test (against 0), respectively. ICC (95% CI) represents ICC(2,1) values and 95% bootstrapped confidence intervals.

|  | **Dorsal**<br>**Dorsal** | **Ventral**<br>**Ventral** | **Within**<br>**Hemicord** | **Between**<br>**Hemicord** |
| --- | --- | --- | --- | --- |
| **Baseline** | r = 0.07<br>t = 12.9<br>p < 0.001<br>ICC (95% CI) =<br>0.71 (0.53 - 0.85) | r = 0.10<br>t = 18.7<br>p < 0.001<br>ICC (95% CI) =<br>0.65 (0.49 - 0.81) | r = -0.01<br>t = -4.7<br>p < 0.001<br>ICC (95% CI) =<br>0.38 (0.22 - 0.54) | r = 0.02<br>t = 7.1<br>p < 0.001<br>ICC (95% CI) =<br>0.28 (0.11 - 0.47) |
| **Baseline +**<br>**WM** | r = 0.06<br>t = 12.9<br>p < 0.001<br>ICC (95% CI) =<br>0.69 (0.51 - 0.84) | r = 0.10<br>t = 18.0<br>p < 0.001<br>ICC (95% CI) =<br>0.66 (0.47 - 0.81) | r = -0.01<br>t = -4.9<br>p < 0.001<br>ICC (95% CI) =<br>0.39 (0.20 - 0.54) | r = 0.02<br>t = 7.2<br>p < 0.001<br>ICC (95% CI) =<br>0.29 (0.10 - 0.47) |
| **Maximal** | r = 0.03<br>t = 9.5<br>p < 0.001<br>ICC (95% CI) =<br>0.59 (0.46 - 0.74) | r = 0.05<br>t = 11.6<br>p < 0.001<br>ICC (95% CI) =<br>0.63 (0.44 - 0.79) | r = -0.02<br>t = -10.7<br>p < 0.001<br>ICC (95% CI) =<br>0.30 (0.06 - 0.53) | r = 0.01<br>t = 6.7<br>p < 0.001<br>ICC (95% CI) =<br>0.18 (-0.03 - 0.38) |
| **Maximal +**<br>**WM** | r = 0.03<br>t = 9.6<br>p < 0.001<br>ICC (95% CI) =<br>0.57 (0.44 - 0.73) | r = 0.05<br>t = 11.5<br>p < 0.001<br>ICC (95% CI) =<br>0.64 (0.46 - 0.80) | r = -0.02<br>t = -11.1<br>p < 0.001<br>ICC (95% CI) =<br>0.29 (0.05 - 0.54) | r = 0.01<br>t = 6.7<br>p < 0.001<br>ICC (95% CI) =<br>0.17 (-0.05 - 0.37) |

## Motivation for estimating correlations in a slice-wise manner

In this study, we calculated correlations for each slice and then averaged the obtained correlations across slices afterwards. While this approach has also been taken by previous studies (e.g. Barry et al., 2014; Conrad et al., 2018), there is also an alternative approach where time series for any given ROI are first averaged across slices and then correlations (between ROIs) computed from these averaged time series (e.g. Eippert et al., 2017; Vahdat et al., 2020). There are two reasons why we did not average the time series from all slices, but rather obtained time series correlations on a slice-wise level and then averaged the resulting correlation coefficients in this study.

The first reason is based on the fact that our field of view (composed of 24 slices) encompasses a large number of spinal segments (C3 to T1) and we assume that the segments represent rather separate functional units based on several lines of evidence. First, Weber et al. (2018) did not find strong evidence for between-segment functional connectivity during rest and describe the observed connectivity patterns as more lattice-like, suggesting that different segments might contain separate networks. Second, Eippert et al. (2017) observed that the similarity of within-segment connectivity patterns decreases with segmental distance. Third, Kong et al. (2014) decomposed resting-state data using spatial ICA and observed segment-like components, whose time series were either uncorrelated or even showed slightly negative correlations. While it might thus be reasonable to assume that time-series in neighboring slices might be similar (especially if they belong to one segment), there is clear evidence to suggest this is not the case across all 24 slices.

The second point we want to raise is best explained by a toy example. Let us assume that we have two regions of interest (e.g. left and right dorsal horns) and acquired time series from these regions in two slices, leading to four time series overall. Let us further assume that the correlation between dorsal horns for each slice is 0.3, so that averaging correlations for each slice would give an overall correlation of 0.3 Let us now assume that the time series from both slices are also correlated with each other, say a correlation of 0.1 from each region of one slice to any region of the other slice. If we create time series that have the described properties and average them before calculating the correlation, we obtain a correlation of 0.36 between the dorsal horns. Thus, depending on the overall correlational structure of the network, the correlation of averaged time series can deviate quite substantially from the average of correlations, as the former is capturing the overall network properties instead of pairwise correlations: importantly, depending on the specific network properties, it can over- or underestimate the slice-wise value. In order to explore this issue in more detail, we refer the reader to a markdown file containing simulations with code and explanations (see associated GitHub repo: https://github.com/eippertlab/restingstate-reliability-spinalcord).

